# PGM3 inhibition Shows cooperative Effects With Erastin inducing Pancreatic cancer cell death via activation of the Unfolded Protein Response

**DOI:** 10.1101/2023.03.19.533311

**Authors:** Barbara Zerbato, Maximilian Gobbi, Tobias Ludwig, Virginia Brancato, Alex Pessina, Luca Brambilla, Andre Wegner, Ferdinando Chiaradonna

**Affiliations:** Tumor Biochemistry, Biotechnology and Biosciences, University of Milano Bicocca, Milan, Italy; Pathometabolism, Department of Bioinformatics and Biochemistry, Braunschweig Integrated Centre of Systems Biology (BRICS), Technische Universität Braunschweig, Braunschweig, Germany.; Center for Genomic Science IIT@SEMM, Italian Institute of Technology, Milan, Italy

**Keywords:** Hexosamine Biosynthetic Pathway_1_, Unfolded Protein Response_2_, Pancreatic cancer cells_3_, Cell Death_4_, Erastin_5_, Ferroptosis_6_.

## Abstract

Pancreatic ductal adenocarcinoma (PDAC) is a highly aggressive cancer with a poor patient prognosis. Remarkably, PDAC is one of the most aggressive and deadly tumor types and is notorious for its resistance to all types of treatment. PDAC resistance is frequently associated with a wide metabolic rewiring and in particular of the glycolytic branch named Hexosamine Biosynthetic Pathway (HBP). Here we show the effect of the combined treatment between an HBP’s Phosphoglucomutase 3 (PGM3) enzyme inhibitor, named FR054, and erastin (ERA), a recognized ferroptosis inducer, on PDAC cell growth and survival. Noteworthy, the combined treatment applied to PDAC cell lines induces a significant decrease in cell proliferation and a concurrent enhancement of cell death. Furthermore, we show that this combined treatment induces Unfolded Protein Response (UPR), NFE2 Like BZIP Transcription Factor 2 (NRF2) activation, a change in cellular redox state, a greater sensitivity to oxidative stress, a major dependence on glutamine metabolism, and finally ferroptosis cell death. Our study discloses that HBP inhibition enhances, through UPR activation, the ERA effect and therefore might be a novel anticancer mechanism to be exploited as PDAC therapy.

## 2 Introduction

Pancreatic ductal adenocarcinoma (PDAC) is a highly aggressive cancer with a poor patient prognosis (1). Conventional therapy, for both resectable and unresectable PDAC relies mainly on the use of the chemotherapeutic agent Gemcitabine (GEM) either alone or in combination with adjuvant therapies such as paclitaxel conjugated to albumin, 5-FU, capecitabine as well as erlotinib which can increase median OS as compared with GEM monotherapy (2). Despite drug combinations improving the median survival rate compared to GEM alone, these novel combinations elicit resistance within weeks and hence fail to bring tumour regression. For this reason, there is an urgent need to search for new pharmacological targets that will boost the sensibility to current treatments and reduce drug resistance.

Tumour metabolic rewiring and metabolic adaptation following drug treatments are typical clues of PDAC (3). Indeed, different reports indicate that reprogrammed metabolism closely regulates PDAC development and chemoresistance (4,5). Among the different metabolic alterations observed in PDAC, the increased flux through the HBP tightly linked with glucose and glutamine metabolism, has been found as a key metabolic change (6,7). Worthy of note, the final metabolite generated by the HBP is the uridine 5′-diphospho-*N*-acetyl-D-glucosamine (UDP-GlcNAc), the main substrate for O- and N-protein glycosylation. Both Post Translational Modifications (PTMs) play critical roles in protein folding, stability, activity, macromolecular interactions, function, and nuclear translocation. Therefore, enhancement of HBP flux, responsible for the aberrant protein glycosylation often observed in different types of tumours including PDAC (8,9) could represent a new target for tumour therapy.

In this regard, we have recently developed and tested in breast and pancreatic cancer cells as well as in vivo models, a novel compound, FR054. This molecule is an N-acetyl-glucosamine 6-phosphate analogue capable to diminish the HBP flux by targeting the HBP enzyme PGM3, leading first to a cell proliferation inhibition and then to cell death (10,11), underlining the fundamental role of this pathway in breast and pancreatic cancer cell proliferation and survival. Importantly, FR054, if combined with GEM (12) or the pan-KRAS inhibitor BI-2852 (13), significantly enhances PDAC cancer cells sensitivity to both drugs, also suggesting a role of HBP in drug resistance.

Our previous findings indicated that the death mechanism induced by FR054 is dependent on the acute activation of the UPR. Indeed, HBP flux reduction, decreasing UDP-GlcNAc availability, causes a reduction in protein N-glycosylation, an accumulation of misfolded proteins, and finally irremediable cell damage, that drives CHOP-dependent apoptotic signalling (11,12). Worthy of note, previous reports propose that the UPR is constitutively active in PDAC and it may contribute to the disease progression and the acquisition of resistance to therapy (14). Thus, these different findings highlight a dual role for UPR in correlation with its level of activation. In particular, it is either an adaptive cellular mechanism to weaken protein and metabolic stresses created by a hypoxic environment and to induce chemoresistance, or it is an apoptotic inducer when prolonged in time (15). For this reason, some authors have suggested that targeting UPR in cancer including in PDAC may be highly beneficial, especially in combinatorial treatments that could provide an effective anti-tumorigenic response in patients.

Therefore, in this study, we aimed to further detail the mechanism of FR054-induced cell death, in order to verify whether its pro-apoptotic ability could be enhanced by co-targeting a specific protein or cellular pathway. For this purpose, we performed an RNA-seq analysis in two different PDAC cell lines, namely MiaPaCa-2 and BxPC3, treated with FR054 to provide unbiased mechanistic insights. Results demonstrate that FR054 treatment induces expression of genes related to UPR, to NRF2 pathway, to glutathione biosynthesis and ferroptosis. Consequently, we show that inhibition of glutathione biosynthesis, by using the specific inhibitor for the Solute Carrier Family 7 Member 11/xCT (SLC7A11), ERA, in combination with FR054, significantly enhanced the FR054 effect causing a noteworthy increase in cancer cell proliferation arrest and death. Remarkable, the latter effect was tightly associated with a higher expression of the pro-apoptotic protein DNA Damage Inducible Transcript 3/CHOP (CHOP), reduced activity of SLC7A11 transporter, an alteration of intracellular glutathione (GSH) and glutathione disulfide (GSSG) levels, and a strengthening of lipid peroxidation indicating that SLC7A11 inhibition is synthetic lethal with FR054. Notably, the metabolic analysis supported the role of FR054 in favouring glutamine metabolism over glycolysis. Therefore, our work reveals that simultaneous inhibition of HBP with FR054 and GSH metabolism could have therapeutic benefits in the treatment of pancreatic cancer.

## 3 Materials and Methods

### Materials

N-Acetyl-L-cysteine (NAC), DL-Buthionine-sulfoximine (BSO), and BPTES were purchased from Sigma-Aldric (Merck Life Science, Milan, Italy), while erastin (ERA) was purchased from Selleckchem (Planegg, Germany). FR054 was synthesized either by our laboratories or by WuXi AppTec Co., Ltd. (Tianjin, China) (10,11).

### Cell lines

Human pancreatic ductal adenocarcinoma cell lines MiaPaCa-2 and PANC1 were routinely cultured in high glucose Dulbecco’s medium Eagle’s medium (DMEM), while BxPC3 cell line in RPMI (Euroclone, Milan, Italy). Both media were supplemented with 2 mM L-glutamine, 100 U/mL penicillin, 100 μg/mL streptomycin (Sigma-Aldrich, Merck Life Science, Milan, Italy) and 10% fetal bovine serum (FBS; Euroclone, Milan, Italy). The cells were grown and maintained according to standard cell culture protocols and kept at 37 °C with 5 % CO2. The medium was replaced every 2-3 days and cells were split or seeded for experiments when they reached the sub-confluence. MiaPaCa-2, PANC1, and BxPC-3 were originally obtained from American Type Culture Collection ATCC.

### Trypan blue vital assay

Where not different specified, for experiments, cells were seeded in the complete growth medium and after 24 h, cells were washed twice with phosphate buffer saline 1X (PBS 1X, Euroclone, Milan, Italy) and incubated in a complete medium with or without FR054. 24 hours later, cells were also treated with or without 10 µM ERA.

Trypan blue vital assay was performed by seeded 4×10^4^ (MiaPaCa-2 and PANC1) and 8×10^4^ (BxPC3) viable cells per well in 12-well dishes, respectively. Inhibition of proliferation and cell death were assessed at different time points (48 and 72 h), counting harvested cells with Burker chamber after staining with trypan blue 0.4% (Life Technologies – Thermo Fisher Scientific, Waltham, MA, USA). MiaPaCa-2 cell line was tested for two different concentrations of FR054, 350 µM or 500 µM, at 72h, and it was decided to use 350 µM as the optimal concentration for all other experiments in all cell lines. Cell death was also measured at 72h of treatments as previously described, using a different treatment scheme: 24 h after seeding, cells were incubated in a complete medium with or without 350 µM FR054, and at 48h post-seeding, cells were also treated with or without 10 µM ERA and 5 mM NAC or 5 mM BSO.

### MTT assay

Cell viability was also assessed with MTT assay (Millipore, Merck Life Science, Germany). Briefly, MiaPaCa-2 and BxPC3 cells were seeded in 96-well plates at the density of 5×10^3^ and 1×10^4^ cells per well, respectively, in the complete growth medium. After 24 h, cells were incubated in a complete medium with or without 350 µM FR054. At 48h post-seeding, cells were also treated with or without 10 µM BPTES or 10 µM ERA. After 48h of FR054 single or combined treatments, tetrazolium salts (0.1 mg/mL) were added directly to the culture medium of each well and incubated for 4 h at 37 °C in a 5% CO2 humidified atmosphere to allow mitochondrial dehydrogenase to convert MTT into insoluble formazan crystals. After incubation, formazan was solubilized by adding 2-propanol 0.1 M HCl (Sigma-Aldrich, Merck Life Science, Milan, Italy). The absorbance was measured at wavelength 570 nm (660 nm background wavelength) with VICTOR X3 multimode plate reader (PerkinElmer, Milan, Italy).

### Western blot analysis

MiaPaCa-2 (6 × 10^5^ cells) and BxPC3 (1,2 × 10^6^ cells) were seeded onto 100 mm dishes in the complete growth medium. After 48 and 72 h, cells were harvested and disrupted in a buffer containing 50 mM Tris-HCl pH 7.6, 150 mM NaCl, 1% (v/v) Triton X-100, 0,2% (v/v) Sodium dodecyl sulphate (SDS), 0,5% (v/v) Sodium deoxycholate, 1 mM MgCl2, 1 mM EDTA, protease inhibitor cocktail, and phosphatase inhibitors (Sigma-Aldrich, Milan); 20 µg of total proteins were resolved by SDS-PAGE and transferred to the nitrocellulose membrane, which was incubated overnight with specific primary antibodies: cleaved caspase-3 (#9662S, 1:500), GRP78 (BiP, #3177, 1:1000), phospho-eIF2α Ser 51 (#3398, 1:1000) from Cell Signaling Technology Inc. (Euroclone, Milan, Italy), eIF2α (sc-133127, 1:1000) from Santa Cruz Biotechnology Inc. (DBA Italia, Milan, Italy), ATF4 (#11815, 1:800), CHOP (#2895, 1:1000) from Cell Signaling Technology Inc. (Euroclone, Milan, Italy), NRF2 (sc-13032, 1:1000) from Santa Cruz Biotechnology Inc. (DBA Italia, Milan, Italy), SLC7A11 (ab175186, 1:5000) from Abcam (Cambridge, UK), HO-1 (HMOX1; #86806, 1:500) from Cell Signaling Technology Inc. (Euroclone, Milan, Italy), Keap1 (sc-365626, 1:1000) and Vinculin (#sc-5573, 1:10000) from Santa Cruz Biotechnology Inc. (DBA Italia, Milan, Italy). Levels of protein expression on Western blots were quantified by densitometry analysis using ImageJ software.

### Glutamic acid and cystine determination

Cells were plated at the density of 4×10^4^ (MiaPaCa-2 and PANC1) and 8×10^4^ (BxPC3) per well in 12-well dishes. The media used for subsequent HPLC analyses were collected after 48h and 72h of treatment. Glutamic acid and cystine concentrations were determined by chromatographic analysis using a Jasco HPLC system (Jasco Europe, Cremella, Lecco, Italy) under the control of the ChromNAV 2.0 software. The analyses were carried out using the method described by Henderson et al. with some modifications. The derivatized amino acids were separated using a Waters (Milford, MA, USA) XTerra RP18 Column (4.6 mm × 250 mm i.d, 5μm particle size) equipped with a precolumn Security Guard (Phenomenex, Macclesfield, UK). The flow rate was set at 1 mL/min with the column oven at 40 °C. Twenty microliters of the sample, appropriately diluted, were injected into the system and the chromatogram was monitored at 338 nm. The pre-column derivatizing reagent o-Phtalaldehyde (OPA) was prepared by dissolving 25 mg of OPA and 25 mg of 3-Mercaptopropionic acid in 5 mL of 0.2 M borate buffer, pH 10.2 and store at 4°C until any sign of degradation. The samples were prepared by adding 5.5 μL of the derivatization reagent to 5.5 μL of the samples and bringing the volume up to 200 μL with water. HPLC separation employed mobile phases A (40mM Sodium Phosphate Buffer, pH 7.8) and B (30% acetonitrile:60% methanol:10% H2O). The column was equilibrated with 94.5%/5.5% (v/v) mobile phase A/B for 10 min before each sample analysis. The elution gradient consisted of 94.5%/5.5% (A/B) for 0.85 min; 2.15-min linear gradient to 13% phase B; 23 min linear gradient to 54% phase B; 94.5%/5.5% (A/B) for 4 min. Amino acids peaks were identified by comparing their retention times with that of the standard mixture. At the end of every set of samples, the column was washed with 20% acetonitrile-80% water for 30 min to remove any trace of salts, and then with 80% acetonitrile-20% water for 20 min and stored in solution.

### Cell Morphology

MiaPaCa-2, PANC1 and BxPC3 cells were plated onto 100 mm dishes in the complete growth medium at the density of 6×10^5^, 6×10^5^ and 1,2×10^6^ per well and they were treated with 350 µM FR054 after 24h. At 48h post-seeding the cells were treated with 10 µM ERA. The images were collected with Olympus CX40 phase contrast microscope using a 20X objective using X_entry Alexasoft Imaging Software.

### GSH and GSSG evaluation

To measure reduced glutathione (GSH) and oxidized glutathione (GSSG) content, MiaPaCa-2, PANC1 and BxPC3 cells were plated in 6-well dishes at the density of 1×10^5^, 1×10^5^ and 2×10^5^ per well, respectively. Intracellular GSH and GSSG levels were determined after 48 h of FR054 and erastin in single or combined treatment by VICTOR X3 multimode plate reader (PerkinElmer, Milan, Italy) using a Total GSH / GSSG colorimetric kit (CO-K097-M, Immunological Sciences, Rome, Italy) according to the manufacturer’s instructions. The sample absorbance was measured at 412 nm. Relative GSH and GSSG content was normalized to the cell number.

### BODIPY assay

To evaluate lipid peroxidation, the BODIPY 581/591 C11 dye (Life Technologies – Thermo Fisher Scientific, Waltham, MA, USA) staining was carried out. One day before treatment, MiaPaCa-2, PANC1 (5 × 10^3^ cells) and BxPC3 (1× 10^4^ cells) were seeded onto a black 96-well plate in the complete growth medium. Then, cells were incubated in the complete medium with or without 350 µM FR054. After 24h, cells were also treated with or without 10 µM ERA. All cell lines were also treated with 5 mM NAC or 5 mM BSO as controls for the presence of oxidative stress. Lipid peroxidation level was measured after 48h of treatment. Cells were washed twice with PBS 1X and incubated in PBS 1X with 10 µM BODIPY C11 dye for 15 minutes at 37 °C. Then, cells were washed again with PBS 1X, and the dye fluorescence was measured. Oxidation of BODIPY C11 resulted in a shift of the fluorescence emission peak from 590 nm to 510 nm proportional to lipid ROS generation and was analyzed using FLOUstar® Omega (BMG Labtech, Germany). The 510/590 ratio was normalized to MTT assay.

### MDA assay

To evaluate the formation of the lipid peroxidation product, malondialdehyde (MDA), the cells were seeded at the density of 6×10^5^ (MiaPaCa-2 and PANC1) or 1,2×10^6^ (BxPC3) onto 100 mm dishes in the complete growth medium. The MDA levels in the cells were measured after 48h of FR054 and ERA in single or combined treatment using Malondialdehyde (MDA) Colorimetric assay kit (CO-K028-M, Immunological Sciences, Rome, Italy) according to the manufacturer’s instructions. The sample absorbance was measured at 532 nm by VICTOR X3 multimode plate reader (PerkinElmer, Milan, Italy) and the relative MDA concentration was normalized to the protein content.

### Stable isotope labeling and GC/MS

All extraction experiments were performed within 3 weeks upon cell thawing. The cells were plated in 6-well dishes at the density of 9×10^4^ cells/well for MiaPaCa-2 and PANC1 cell lines or 1.1×10^5^ cells/well for BxPC3 cell line. After 24 hours of seeding, the media were replaced with DMEM for Mass Spectrometry (Sigma-Aldrich, Merck Life Science, Milan, Italy) for MiaPaCa-2 and PANC1 cell lines, and with MS-Grade SILAC RPMI 1640 (Gibco, Thermo Fisher Scientific, Waltham, MA, USA) medium, without Phenol Red, L-Arginine, and L-Lysine which were manually added at a concentration of 200 mg/mL and 40 mg/mL respectively, for BxPC3 cell line. In both media, it was added 10% sterile-filtered dialyzed FBS (Gibco, Thermo Fisher Scientific, Waltham, MA, USA). For stable isotope labeling, in media, U13C Glucose and unlabeled Glutamine or U13C Glutamine and unlabeled Glucose (Sigma-Aldrich, Merck Life Science, Milan, Italy) were also added. After 48h of FR054 and ERA in single or combined treatment, the medium of each condition was collected, and wells were washed with 0.9% NaCl solution to perform metabolite extraction following standard laboratory procedure as described in (16). Then, the polar phase (upper phase) was collected, dissolved in MeOH/IS water, and transferred to GC/MS glass vial. The eluates were evaporated in a vacuum concentrator system (CentriVap; Labconco, Kansas City, MO, USA) and stored at -20°C until further analysis. Targeted analysis was carried out on Agilent 7890B Gas Chromatograph equipped with a 30-meter DB-35MS + 5-meter Duraguard capillary column. Helium was used as carrier gas at a flow rate of 1 mL per minute. The system was connected to an Agilent 5975 Mass Spectrometer (Agilent Technologies, Santa Clara, CA, USA). Derivatization reagent N-tert-Butyldimethylsilyl-N-methyltrifluoroacetamide, 1 % tert-Butyldimethylchlorosilane (MTBSTFA with 1 % t-BDMCS; Restek Corporation, Bellefonte, PA, USA) and Methoxylamine (MeOX), dissolved in pyridine at a concentration of 20mg/mL were added to protect metabolites. Mixing was carried out by an autosampler dissolving in MeOX/pyridine and shaking at 40°C for 90 minutes. Afterward, derivatization agents were injected into vials, shaken for 30 or 60 minutes at 40 °C, and analyzed. GC oven was set to 100 °C for 2 minutes, increasing temperature 10 °C per minute until it reached 300 °C. Temperature was maintained for 3 minutes, raising it then another 25 °C. MS performed the electron ionization at 70 eV, the source had a constant temperature of 230 °C, and quadrupoles had a constant temperature of 150 °C. Analytes were detected in Selected Ion Monitoring (SIM) mode. Raw GC/MS data were analyzed on MetaboliteDetector, an open-source software running on Linux-based operative systems developed by Karsten Hiller et al.(17). Chemical formulas for Mass Isotopomer Distribution (MID) determination were taken from (18).

### Gene expression analysis

Two biological replicates were used for gene expression analysis. In particular, MiaPaCa-2 and BxPC3 cell lines were seeded at the density of 5×10^6^ cells in 100 mm dishes and after 24h were treated with 350µM FR054 and incubated for 48h. Then, the cells, untreated and treated, were pelletized and the mRNA was extracted by using Qiagen RNeasy Kit (X) starting from 5×10^6^ cells/samples. RNA-seq experiments, data extraction, and analysis were performed in outsourcing (GALSEQ, Milan, Italy). In particular, total RNA was quantified with a spectrophotometer method and RNA integrity was evaluated with Agilent 4200 TapeStation analysis. RNA stranded libraries were generated starting from 500ng total RNA and mRNA was selected with poly-T oligo attached magnetic beads using Illumina TruSeq® Stranded mRNA Library Prep kit (Catalog #: 20020594). Libraries were quantified and quality checked on an Agilent 4200 TapeStation system (High Sensitivity D1000), and then sequenced on an Illumina NovaSeq 6000 platform with paired-end reads 150bp long, with a depth of 30 million cluster/sample. Fastq reads were quality checked using FastQC (https://www.bioinformatics.babraham.ac.uk/projects/fastqc/) for overall and per-base read quality and presence of adaptor sequences. Paired reads were subsequently aligned to the human reference genome (GRCh38/hg38) using the splice-aware aligner STAR v 2.5.0c and indexed with Samtools. Per-gene read counts were generated using the STAR quantMode geneCounts option. Raw per-gene read counts were then processed using the Bioconductor package DESeq2 package v. 1.30 (19) in order to perform the differential expression analysis. Genes with an adjusted p-value (Benjamini-Hochberg False Discovery Rate - FDR) < 0.05 and log2FC <-2 or ≪2 were considered as differentially expressed. Sorted, indexed bam files were finally used for manual quality check of the alignment profiles. The RNA-seq data have been deposited at NCB-GEO database, accession number GSE223303.

### Statistical analysis

All statistical analyses were performed using GraphPad Prism v 8.0.2 (GraphPad Software Inc., La Jolla, CA) and data are presented as mean ± SD and as mean ±SEM from three or more independent experiments. For the inhibition of proliferation evaluated after treatment with different concentrations of FR054 and/or ERA and for GSH, GSSG and MDA determination, statistical significance (*p < 0.05, **p < 0.01, ***p < 0.001 ****p < 0.0001) was determined using one-way ANOVA with Tukey’s multiple comparisons test. For GC/MS analysis performed with stable isotope labeling statistical significance (*p < 0.05, **p < 0.01, ***p < 0.001 was performed with Students T-Test. Instead, for all other experiments, statistical significance (*p < 0.05, **p < 0.01, ***p < 0.001 ****p < 0.0001) was determined using two-way repeated measures ANOVA with Tukey’s multiple comparisons test.

## 4 Results

### Transcriptional analysis reveals upregulation of NRF2 and ferroptosis pathways following PDAC cells treatment with FR054

To gain insights into the transcriptional differences between KRAS PDAC mutated cell line, MiaPaCa-2, and wild-type PDAC cell line, BxPC3, upon FR054 treatment, we compared FR054-regulated transcriptome to untreated samples in MiaPaCa-2 and BxPC3 cells by RNA-seq analyses. To find differentially expressed genes, we used mRNAs showing 2-fold changes with p-values smaller than 0.05. The analysis captured in MiaPaCa-2 cells, 1648 differentially expressed genes (DEGs), among which 910 were significantly upregulated and 738 significantly downregulated in FR054-treated cells as compared to untreated cells (Figure 1A, Supplementary Table 1). In BxPC3 cells the analysis identified 373 DEGs among which 100 were significantly upregulated and 273 significantly downregulated in FR054-treated cells as compared to untreated cells (Figure 1B). We found that genes containing antioxidant response element (ARE) regulated by NRF2 protein, including *Heme Oxygenase 1 (HMOX1), Glutamate-Cysteine Ligase Catalytic Subunit (GCLC), Glutamate-Cysteine Ligase Modifier Subunit (GCLM), Spermidine/Spermine N1-acetyltransferase 1* (*SAT1*), *Aldo-Keto Reductase Family 1 Member C1* (*AKR1C1*), *Oxidative Stress Induced Growth Inhibitor 1 (OSGIN1), Glutathione-Disulfide Reductase (GSR), SLC7A11, Thioredoxin Reductase 1 (TXNRD1) Ferritin Light Chain (FTL), Activating Transcription Factor 3 (ATF3), Aldehyde dehydrogenase 3A1 (ALDH3A1), Alpha/beta-hydrolase domain containing 4 (ABHD4)* were the top enriched genes in treated cell lines compared with untreated cells. Worthy of note also the other top enriched genes were related to the anti-oxidant response regulated by NRF2 since *MAF BZIP Transcription Factor G* (*MafG*) heterodimerizes with NRF2 and *NmrA Like Redox Sensor 1* (*NMRAL1P1*), which encodes for an NADPH sensor protein, is tightly associated with NRF2 pathway. In addition, as expected for the well-known capacity of FR054 to induce ER stress and UPR, some transcription factors, tightly linked with this response, including *DNA Damage Inducible Transcript 3/CHOP (DDIT3), Early Growth Response 1 (EGR1) and Activating Transcription Factor 3 (ATF3)* were more upregulated in MiaPaCa-2 treated cells. To computationally confirm the specific enrichment of some transcription factors (TFs) as responsible for the observed changes in gene expression upon FR054 treatment, up-regulated DEGs for both cell lines were used as query to interrogate the ChEA3-ChIP-X Enrichment analysis libraries using the web-based software Enrichr (20). In particular we used for MiaPaCa-2 cells 700 out 910 upregulated DEGs with a log^2^FC *≥* 2 and the ReMap library (21). Conversely, since the upregulated DEGs were of low number for BxPC3 cells, specifically 100 but only 80 were recognized by the database, we decided to accept for the transcription factors enrichment analysis also the upregulated DEGs with a log^2^FC *≥*1.5 (upregulated genes used: 137) and the Top rank analysis that gives as readout the integrated results from all the ChEA3-ChIP-X libraries. As shown in Supplementary Table 2 and 3, in MiaPaCa-2 cells, among the top ten enriched TFs, were identified not only NRF2 (NFE2L2) but also other TFs related to NRF2 activity including CEBPB, ATF3, and MAFF. Equally in BxPC3 cells, we identified NRF2 and some other TFs related to NRF2 activity including MAFG, ATF3 and BACH1. In line with these results, gene set enrichment analysis (GSEA) of the RNA-seq transcriptional profiling showed in both cell lines an upregulation of NRF2, UPR, and ferroptotic pathways in treated cells compared with control cells (Figure 1C, D, Figure S1, S2). To further detail the activation of NRF2 and ferroptosis related genes, we searched in our RNA-seq data, mRNAs (p-values smaller than 0.05) related to NRF2-dependent activation mechanism as well as genes related to ferroptosis. In particular, we identified six different cellular processes linked to NRF2 activity and/or ferroptosis namely gluthatione metabolism, iron metabolism, ROS metabolism, lipid metabolism, metabolism and autophagy as well as transcription factors associated with NRF2. As shown in the dendrograms presented in Figure 2A, several mRNAs linked to these selected processes were significantly regulated upon FR054 treatment, especially in MiaPaCa-2 cells compared to BxPC3 cells. Indeed FR054 treatment, transcriptionally upregulated most of the genes involved in GSH metabolism, *cystine/glutamate antiporters SLC7A11* and *SLC3A2* and *Glutathione-Disulfide Reductase (GSR)*, glutamine, cysteine and glycine metabolism (*GLS1, GCLM, GCLC, GOT1, PHGDH, PSAT1*), NADPH production (*G6PD, PGD, ME1*) as well as genes involved in anti-ferroptosis mechanisms such as iron transport and storage (*FTH1, FTL*), Fe2^+^ sequestration (*NUPR1* and *LCN2*), membrane repair mechanisms (*AIFM2, NQO1, AKR1Cs*) and autophagic genes associated with both NRF2 activation and ferroptosis (*GABARAPL1, MAP1LC3B2, SQSTM1*). Conversely, genes associated with lipid metabolism, generally repressed by NRF2, such as *ACACA, SCD, LPCAT3, FADS2*, and *ELOVL6* were downregulated, further confirming an FR054-dependent activation of the NRF2 axis. The latter effect was also confirmed by the concomitant upregulation of different transcription factors related to the NRF2 pathway including, *MAFA, MAFG, MAFF, CEBPB, ATF4*, and *NRF2* as well. Of note, FR054 induced also a few important ferroptosis genes including *CHAC1*, a γ-glutamyl cyclotransferase toward glutathione, a well-known ferroptosis marker, and HMOX1, the major intracellular source of iron Fe^2+^ (Figure 2B). Several of these genes were equally regulated in BxPC3 but to a lesser extent suggesting a lower antioxidant response as compared to MiaPaCa-2 cells upon FR054 treatment. Altogether, transcriptional data demonstrate that, in both cell lines, FR054 treatment is associated with the induction of genes involved in a large NRF2-dependent antioxidant response and a significant alteration of genes associated with either inhibition or induction of ferroptosis (Figure 2B), and therefore suggesting that drugs interfering with these mechanisms could be used in the experiment of synthetic lethality in combination with FR054.

**Figure 1.**
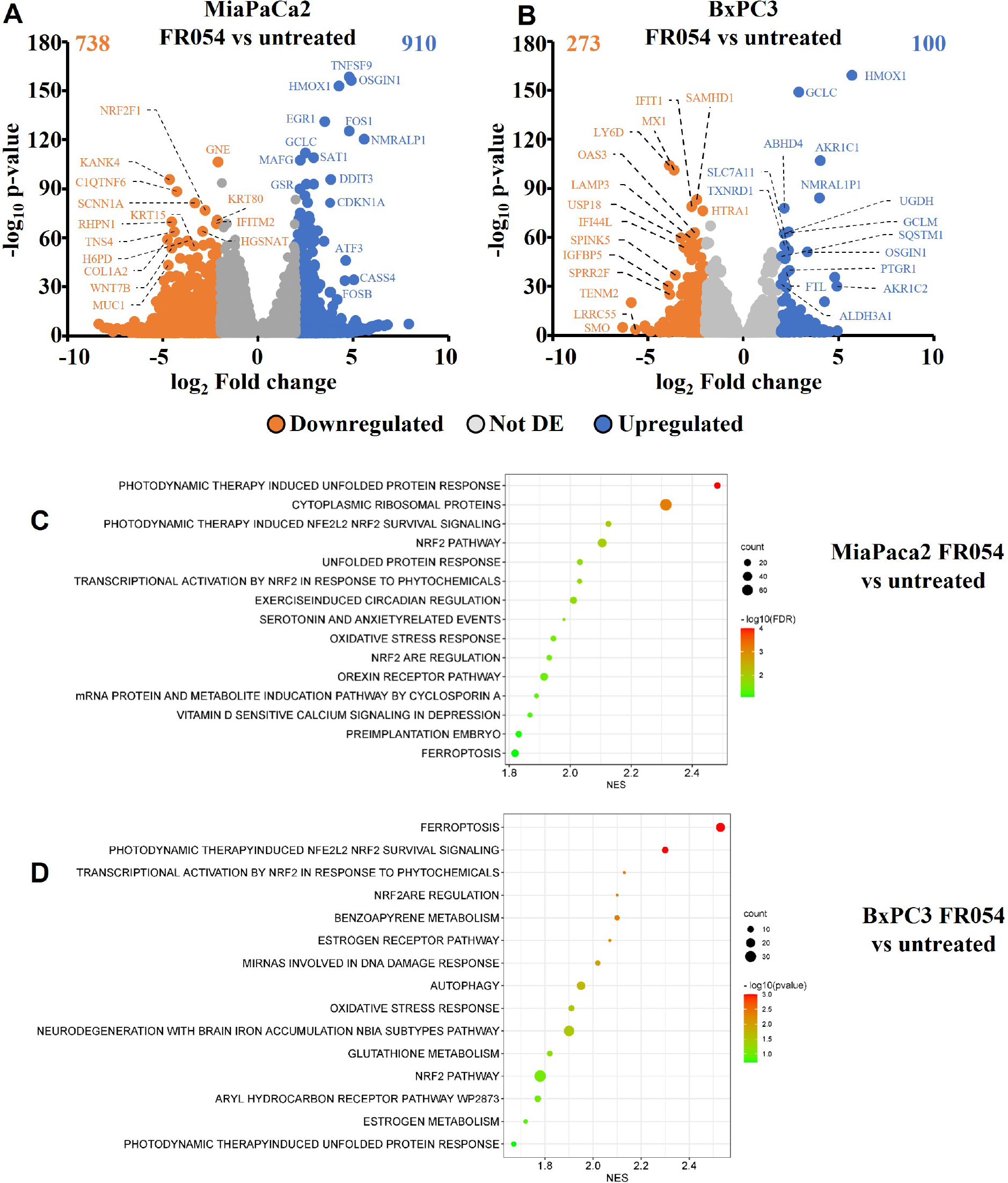
Identification and characterization of DEGs in PDAC cell lines upon FR054 treatment compared to control. **(A) and (B)** Volcano plot of DEGs identified in MiaPaCa-2 and BxPC3 FR054 treated cells versus untreated cells. Upregulated genes are indicated with cyan dots; downregulated genes are indicated with orange dots; not DEGs are indicated with grey dots. DEGs p-value <0.05. Top 15 DEGs are depicted; **(C) and (D)** Gene set enrichment analysis (GSEA) performed by using the DEGs shown in A and B of MiaPaCa-2 and BxPC3 FR054 treated cells versus untreated cells.

**Figure 2.**
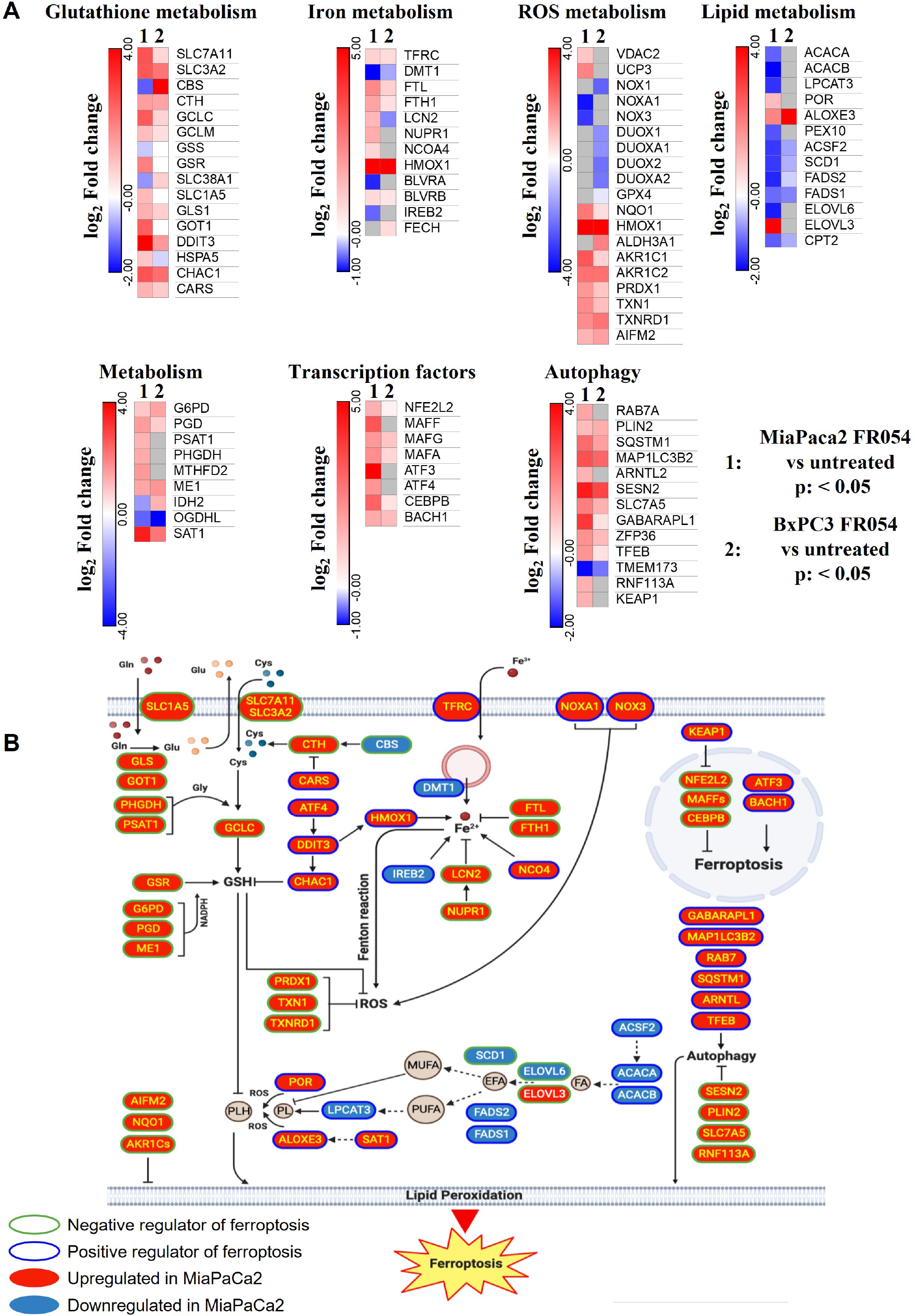
DEGs in MiaPaCa-2 and BxPC3 FR054 treated cells versus untreated cells are enriched of mRNAs involved in oxidative stress response and ferroptosis. **(A)** Dendrograms show genes belonging to the cellular processes described in the figure that are present in the list of the DEGs (p-value <0.05, Benjamini-Hochberg False Discovery Rate - FDR). In the heat map color indicated log^2^ fold change, red represents upregulated genes, blue represent downregulated genes, white no DEGs, and grey missing values; (B) The cartoon shows the role of some of the genes identified in (A) in ferroptosis process as described in the main text.

### Inhibition of glutamine/glutamate metabolism enhances the effect of FR054 by increasing PDAC cell death

NRF2 is a master regulator of antioxidative responses and plays critical roles in maintaining redox balancing so NRF2 activators attenuate oxidative stress. Indeed, NRF2 is one of the key regulators of GSH metabolism. GSH synthesis enzyme glutamate-cysteine ligase, glutathione synthetase, and reductase are target genes of NRF2 (22). Glutaminase (GLS), which can catalyze glutamine into glutamate is also promoted by NRF2 (23). Besides, the expression of SLC7A11, the light chain subunit of the Xc-antiporter system (xCT) is activated by NRF2. xCT, a translocator of cystine and glutamate, imports and exports cystine and glutamate into and out of the cell, respectively. Since our transcriptional data clearly indicated an activation of the GSH metabolism upon FR054, we tested if inhibition of glutamine/glutamate utilization by PDAC cells could increase FR054 cytotoxicity, enlightening a cooperative lethal effect between HBP inhibition and GSH synthesis inhibition. To this end, MiaPaCa-2 cells were first treated for 24h with FR054 (350μM) and then treated for a further 24h with two well-known inhibitors, ERA (10μM) that blocks the activity of the antiporter SLC7A11 and Bis-2-(5-phenylacetamido-1,3,4-thiadiazol-2-ylethyl sulfide, -BPTES- (10μM) that blocks the activity of GLS1, either alone or in combination with FR054 (Figure S3A for the experimental scheme).

In MiaPaCa-2 cells, as result of vital cell count, ERA treatment reduced cell proliferation of around 30%, BPTES of around 43% and FR054 of around 52% compared to untreated cells (Figure 3A), suggesting an important proliferative role of glutamine metabolism in this cell line. Remarkably, the FR054 anti-proliferative effect was significantly enhanced when combined with ERA and BPTES since the proliferation reduction reached the values of 69% and 61% in combined treatments, respectively.

**Figure 3.**
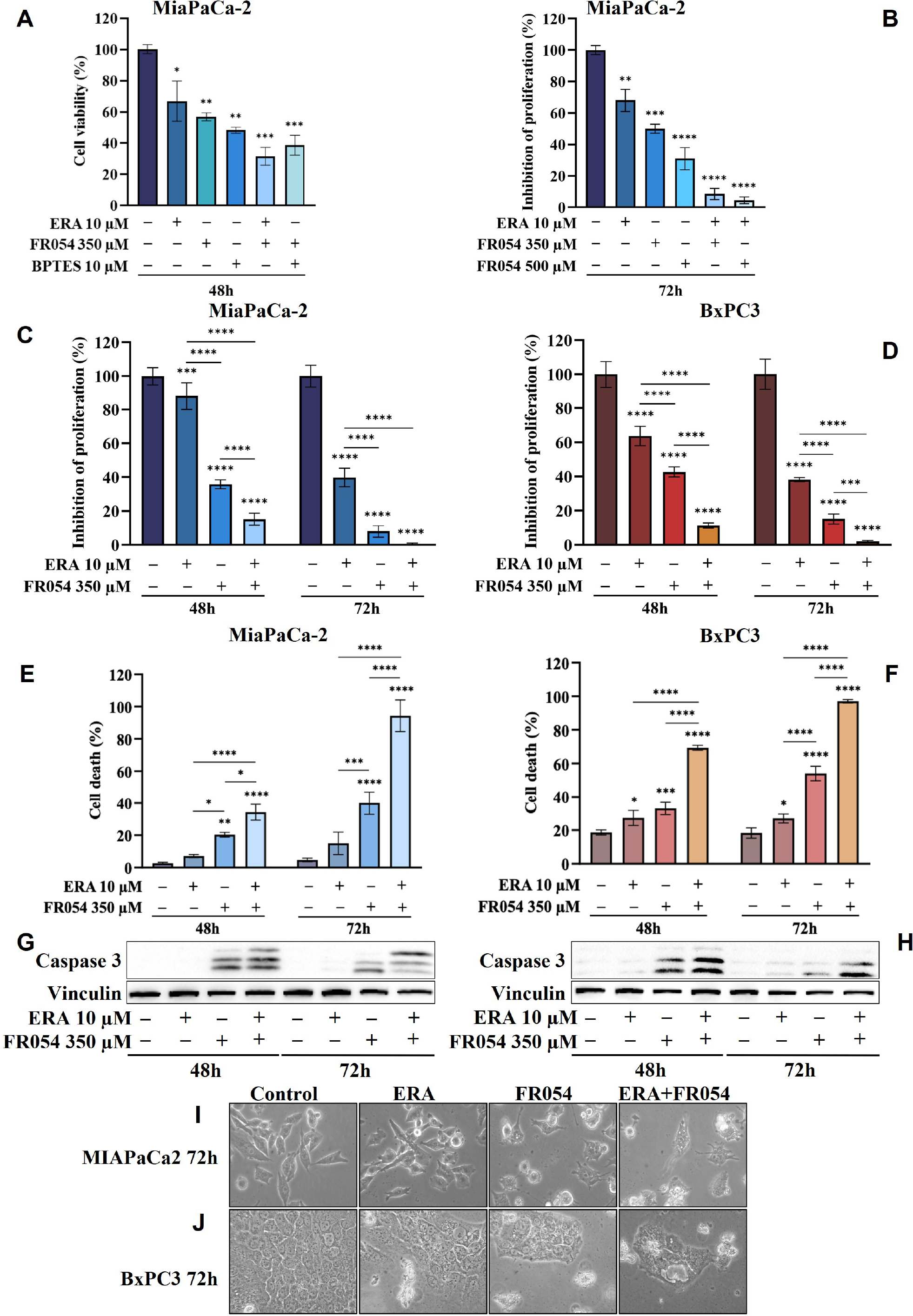
Combined treatment with FR054 and erastin enhances cell proliferation arrest and death of PDAC cell lines. **(A)** Trypan blue cell count after 48h of treatment with FR054, erastin (ERA) 24h, BPTES 24h, or the combination of FR054 with ERA or BPTES added after 24h of FR054 treatment; **(B)** Inhibition of proliferation expressed as percentage (trypan blue assay) after 72h of treatment with two different doses of FR054 alone or in combination with ERA; **(C)** and **(D)** Inhibition of proliferation expressed as percentage (trypan blue assay) in MiaPaCa-2 and BxPC3 cells after 48 and 72h of treatment with FR054 and ERA only or in combination; **(E)** and **(F)** Death cell percentage (trypan blue assay) in MiaPaCa-2 and BxPC3 cells after 48 and 72h of treatment with FR054 and ERA only or in combination; **(G)** and **(H)** Western blot analysis of caspase 3 cleavage in cells treated as (C) and (D). Vinculine has been used as internal loading control; **(I)** and **(J)** Bright-field images of MiaPaCa-2 and BxPC3 cells treated as (C) and (D) taken 48h after the treatments. *p < 0.05, **p < 0.01 ***p < 0.001, ****p < 0.0001. The data are presented as mean ± SD from three independent experiments.

Taking into consideration the greater outcome of the combination of FR054 and ERA, we decided to further detail this effect by increasing the amount and the time of the treatment with FR054. In particular, the cells were treated as described in the scheme of Figure S3B. Briefly, cells were treated for 24h with two different amounts of FR054 (350 and 500μM) and then treated for a further 48h with ERA (10μM) alone or in combination with FR054. The cell survival was analyzed by counting the cells at 72h. As shown in Figure 3B, 48 hours of ERA treatment caused almost a 30% reduction in cell proliferation. As previously published, a dose-dependent cell proliferation arrest was observed in MiaPaCa-2 cells treated for 72 hours with FR054 (12). Indeed, FR054 treatment at 350 and 500μM caused a cell number reduction of approximately 36% and 70%, respectively, as compared to control cells (Figure 3B). Remarkably, such an anti-proliferative effect was significantly enhanced by ERA since the proliferation reduction reached 92% and 95% at 72h in combined treatments, suggesting that SLC7A11 antiporter activity is necessary for cancer survival under HBP inhibition.

To further evaluate the cooperative effect of ERA when combined with FR054, we tested the combination in MiaPaCa-2 and BxPC3 cells along a time course of 72h, by using the lowest dose of FR054 (350 μM) and analysing both proliferation and cell death by vital trypan blue staining. As shown in Figures 3C and D, ERA alone at 48h, as previously published (24), had a lower effect in MiaPaCa-2 cells as compared to BxPC3. Nevertheless, at 72h, in both cell lines induced almost 60% of cell proliferation arrest. FR054 had a stronger effect as compared to ERA, at both time points and in both cell lines. Strikingly, the combined treatment at 72h induced around 100% of cell proliferation reduction, confirming the significant effect of this combination. The cytotoxic analysis additionally confirmed the above results (Figure 3E, F). 24h and 48h of ERA treatment, in both cell lines, induced a little effect on cell death. Conversely, FR054 treatment (48h and 72h of treatment) was significantly more cytotoxic since cell death values were significantly higher at both time points. Strikingly, combined treatment, significantly enhanced cell death in both cell lines at both time points causing almost 100% cell death at 72h (Figure 3E, F), strongly confirming the effective cooperative outcome between FR054 and ERA in PDAC cells. As ERA is known for its pro-ferroptosis activity (25) and FR054 was shown to be able in inducing apoptosis (11,12), next we analyzed the activation/cleavage of caspase-3 as an exclusive apoptotic marker. At 48 and 72h in FR054 treated cells as well as in combined treatment, caspase-3 signals increased as compared to control. However, despite the significant increase in cell death observed at 72h (Figure 3C, D), caspase-3 cleavage was less pronounced as compared to 48h, suggesting also a non-apoptotic mechanism of cell death (Figure 3G, H, Figure S4, S5). Conversely, in ERA alone at both time points, despite the increased cell death observed in particular in BxPC3 cells (Figure 3C, D), caspase-3 signal was not or barely detectable, confirming that the ERA was inducing a non-apoptotic cell death (Figure 3G, H, Figure S4, S5). Besides, cell morphological study further confirmed that combined treatment induced different morphological alterations as compared to single treatments (Figure 3I, J). Altogether these findings support the notion that the combined treatment induces both an apoptotic and a non-apoptotic cell death.

### Cell death dependent on FR054 and ERA combination is associated with ER stress persistence and NRF2-dependent antioxidative response inactivation

Previously published data indicated that prolonged treatment with FR054 causes cell death by inducing ER stress and activation of UPR (11,12). On the other hand, ERA treatment induces cysteine depletion and oxidative cell stress, causing both ER stress and then ferroptosis (26). Furthermore, transcriptional data shown in Figure 2A, further suggest activation of UPR and of antioxidant response, especially in MiaPaCa-2 cells. Therefore, we decided to evaluate UPR activation and cellular oxidative stress upon single and combined treatments by immunoblot analysis.

For UPR activation, we focused our analyses on eIF2α phosphorylation and protein expression of GRP78, linked to ER stress, ATF4, linked both to ER stress and cysteine depletion (27) and CHOP, a transcription factor associated with cell cycle arrest (28) and induction of pro-apoptotic signals upon prolonged UPR activation (29). ERA treatment of MiaPaCa-2 cells, at both 48 and 72h, resulted in an increased expression of GRP78, ATF4, and CHOP as compared to control cells (Figure 4A, Figure S4). No significant change in eIF2α phosphorylation was observed. FR054 treatment resulted in an increase of eIF2α phosphorylation, a slight increase of ATF4 at both 48h and 72h, and a more significant increase of CHOP as compared to control cells. No significant change in GRP78 expression was observed (Figure 4A, Figure S4). Of note in FR054-treated cells, ATF4 was induced at significantly lesser extent as compared to ERA treatment. Combined treatment resulted in higher levels of eIF2α phosphorylation, GRP78 and CHOP as compared to control and single drug-treated cells. ATF4 was induced at 48h to significantly decrease at 72h. Nevertheless, ATF4 level was significantly lower as compared to ERA-treated cells (Figure 4A, Figure S4). Next, we analyzed the behavior of the same proteins in BxPC3 cells, characterized by the expression of a wild-type K-Ras and known to be more sensitive to ferroptosis inducers. As shown in Figure 4B and Figure S5, in BxPC3 cells treated with ERA, the different UPR markers were generally upregulated but in no significant manner (eIF2α phosphorylation, ATF4, and CHOP) or downregulated at significant manner (GRP78). FR054 treatment resulted in a time-dependent increase of eIF2α phosphorylation and CHOP, in a not significant increase of ATF4 and a significant downregulation of GRP78 at 48h (Figure 4A, Figure S4). Conversely, in cells treated with the combination, the three UPR markers, eIF2α phosphorylation, GRP78 and CHOP, increased significantly in a time-depended manner, especially at 72h, as compared to control and single-treated cells (Figure 4B and Figure S5). Of note in combined treatment, ATF4 expression, conversely to what observed in MiaPaCa-2 cells, increased significantly only at 48h to decrease at 72h. Taking together, our findings, in line with previous reports showing the protective role of ATF4 (30,31), could suggest that the ERA-resistance of MiaPaCa-2 cells as compared to BxPC3 cells may be ascribed to a higher ATF4 level of expression as compared to BxPC3 cells, and that, such resistance is hindered by the ability of combined treatment to induce UPR, as highlighted by increased eIF2α phosphorylation as well as GRP78 and CHOP higher expression, in tight association with ATF4 decrease. Indeed, in the combination-treated cells, the sample in which we observed the highest level of cell death (Figure 3C), CHOP remained always higher as compared to ATF4. Such a protective role of ATF4 is further suggested by BxPC3 results, in which the highest values of cell death observed in combination-treated cells (Figure 3D, 72h) were associated with low and high levels of ATF4 and CHOP, respectively. Previous transcriptional data indicated that FR054 treatment can transcriptionally induce an NFR2-dependent antioxidant response, proposing that FR054, as hitherto shown in breast cancer cells (11), induces cellular oxidative stress in PDAC cells. Since several data indicated that NRF2 and ATF4 interact and cooperatively upregulate a number of antioxidant and antiapoptotic genes involved in a protective response engaged during ER stress (32,33) we decided to determine NRF2 expression under our experimental conditions. As expected, upon 48h, the protein level of NRF2 was upregulated in response to ERA and FR054 and such upregulation was stronger in combined treatment in both cell lines (Figure 4A, B, Figure S4 and S5), indicating increased oxidative stress. Surprisingly, at 72h, while in ERA-treated cells, NRF2 expression remained high (MiaPaCa-2 cells) or further induced (BxPC3 cells), conversely in FR054-treated samples, especially upon combined treatment, it was significantly reduced. This behavior was observed especially in ERA-resistant MiaPaCa-2 cells further suggesting that ATF4 and NRF2 levels may dictate ERA resistance (Figure 4A, B and Figure S4 and S5). Oxidative stress as well as UPR, stabilizing and activating NRF2, regulates downstream gene transcription, including SLC7A11 and HMOX1 (34–36). Former, amplifies glutamate secretion, cystine uptake, and facilitates GSH synthesis for Reactive Oxygen Species (ROS) detoxification (37), latter helps keep the cellular redox balance by catalyzing the degradation of heme to carbon monoxide, iron, and biliverdin (38). Thus, to demonstrate NRF2 activation, we analyzed SLC7A11 and HMOX1 protein levels. As shown in Figure 4A and Figure S4, in MiaPaCa-2 cells both SLC7A11 and HMOX-1 almost mirrored NRF2 expression at both time points. Conversely, in BxPC3 cells, both proteins were less modulated and only partially mirrored NRF2 expression, since both were only upregulated at 48h and in particular SLC7A11 slightly increased only in ERA-treated cells and HMOX1 only in combined-treated cells. Nevertheless, both proteins significantly decreased at 72h (Figure 4B, Figure S4 and S5), further suggesting that the major sensitivity of BxPC3 cells to ERA and combined treatments may be ascribed to low ATF4 expression and NRF2 activation. To explore the oscillatory expression of NRF2 protein upon single and combined treatment, then we determined the expression level of *Kelch-like ECH-associated protein 1* (KEAP1), a well-known negative regulator of NRF2 protein stability (39). The expression of KEAP1 significantly increased only in FR054-treated cells and at 48h since at a later timepoint the level decreased, especially in combined treatment (Figure 4A and Figure S4 and S5), therefore justifying the reduced NRF2 expression in these cell samples. Altogether these data suggest that increased cell death in combined treatment (Figure 3E, F), is tightly linked to a significantly higher expression of the pro-apoptotic protein CHOP, a greater antioxidative response at an early time point, as confirmed by the higher NRF2, SLC7A11 and HMOX1 levels, that at later time point, together with ATF4, abruptly decrease.

**Figure 4.**
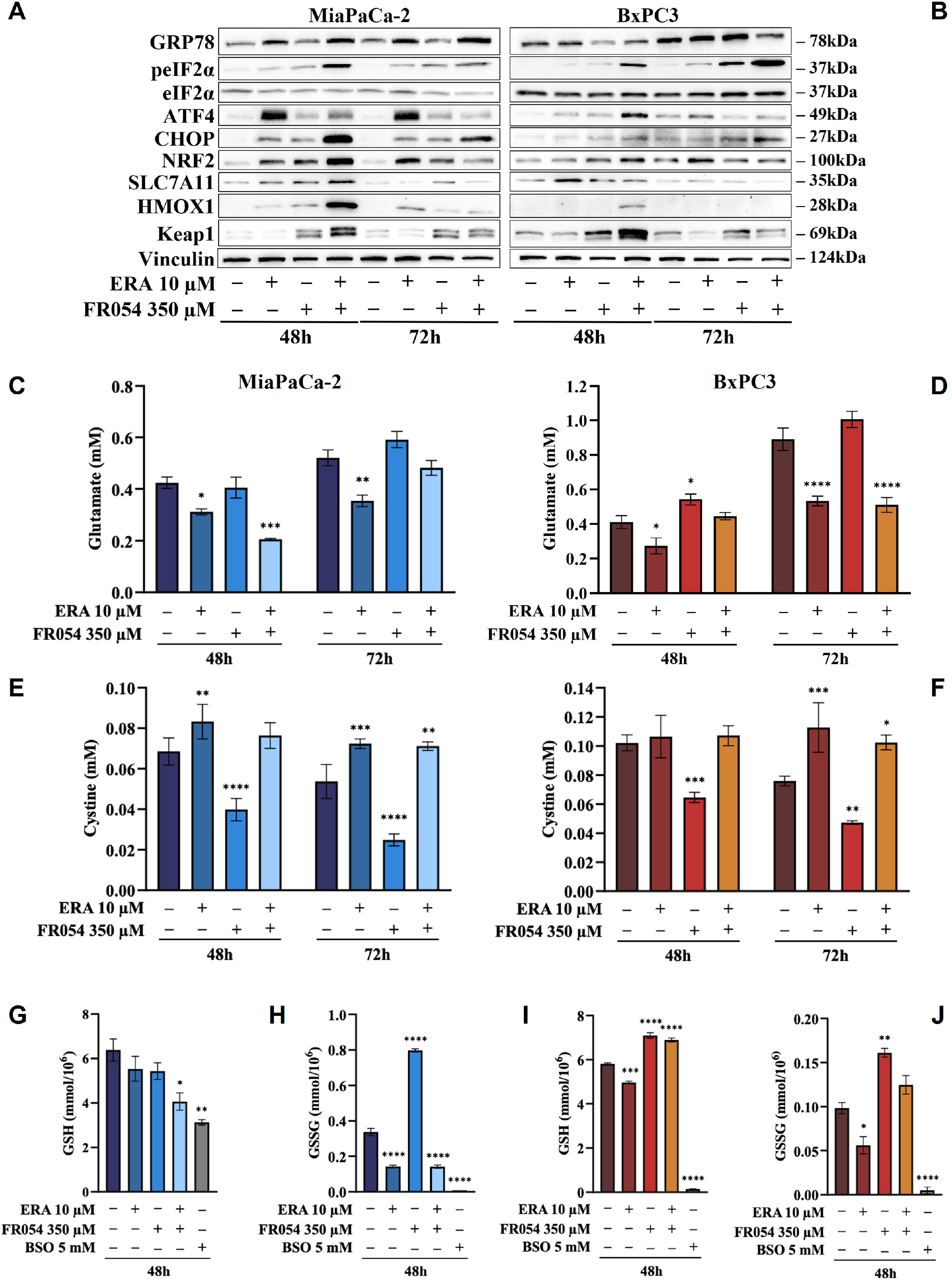
FR054 alone or in combination with erastin affects UPR and oxidative response proteins expression, modifies SLC7A11 activity and alters GSH and GSSG intracellular levels. **(A)** and **(B)** Immunoblot analysis showing the effects of the two compounds on different UPR and oxidative stress response proteins; **(C)** and **(D)** HPLC evaluation of the glutamate concentration in the growth medium of the MiaPaCa-2 and BxPC3 cells after the different treatments as indicated in figure; **(E)** and **(F)** HPLC evaluation of the cystine concentration in the growth medium of the MiaPaCa-2 and BxPC3 cells after the different treatments as indicated in figure; **(G)** and **(H)** measurement of the intracellular concentration of GSH (I) and (J) and GSSG in MiaPaCa-2 and BxPC3 cells after the different treatments as indicated in figure. *p < 0.05, **p < 0.01 ***p < 0.001, ****p < 0.0001. The data of GSH and GSSG concentration are presented as mean ± SEM from three independent experiments, while the other ones are presented as mean ± SD from three independent experiments

### FR054 modulates SLC7A11 function without restraining the inhibitory effect of erastin

To better understand the relation between the observed level of SLC7A11 protein and its function as an exchanger of the anionic form of cystine and glutamate, we measured the amount of these two amino acids into cultured media in both cell lines. While ERA, as expected, inhibited glutamate release in both cell lines and at both time points analyzed, FR054 alone, as predictable by the transcriptional data, albeit not statistically significant, caused a time-dependent increase in the amount of released glutamate (Figure 4C, D). Of note, despite the different temporal dynamics, ERA combined with FR054 still inhibited glutamate release in both cell lines counteracting the positive effect of FR054 treatment (Figure 4C, D). Since glutamate is secreted in exchange with cystine uptake, necessary for GSH synthesis, then we analyzed extracellular levels of cystine. Both lines treated with ERA alone or in combination with FR054 showed a cystine uptake reduction as compared to control cells, mirroring the effect on glutamate efflux (Figure 4E, F). Conversely, in FR054-treated cells a significant reduction of extracellular cystine was observed, confirming the ability of FR054 to increase the exchange between glutamate and cystine (Figure 4E, F). The latter result was confirmed also in BxPC3 cells treated with FR054 alone given that also in this cell line there was a significant decrease of extracellular cystine (Figure 4D). Given that one molecule of cystine can then be converted into two molecules of cysteine, which is a committed step for GSH biosynthesis next, we evaluated at 48h the intracellular level of GSH and GSSG. As technical control, we treated the cells for 24h with 5 mM buthionine sulphoximine (BSO) alone, an inhibitor of GCLC activity. GSH in MiaPaCa-2 cells was statistically significantly reduced only in cells treated with the combination of the two drugs as compared to control cells. Indeed, in ERA and FR054 treated cells, GSH still decreased but not in a statistically significant value (Figure 4G). Conversely, a significant variation of GSSG levels was observed in all treated cells as compared to the control (Figure 4H). In fact, in cells treated with ERA alone or in combination with FR054, accordingly to the significant reduction of cystine uptake, also the GSSG levels were drastically reduced (Figure 4H). A similar behavior was observed in BSO-treated cells, in which the prolonged inhibition of GCLC caused a significant decrease of both GSH and GSSG, as previously observed (40). Surprisingly, in FR054 treated cells, consequently to the significant cystine uptake, there was a significant accumulation of GSSG (Figure 4H). Analysis of GSH and GSSG levels in BxPC3 as compared with the control group showed that ERA induced a slight but significant reduction in both GSH and GSSG levels (Figure 4I, J). Conversely, in FR054 treated cells, either alone or in combination with ERA, both GSH and GSSG levels remained higher as compared to control cells (Figure 4I, J). Of note BxPC3 cells were more sensitive to BSO treatment since 24h treatment completely depleted both GSH and GSSG. Such a result in combined treatment was in part unexpected because following this treatment a significant reduction in extracellular glutamate as well as in cystine uptake was observed. However, based on our transcriptional data (Figure 2A), we cannot exclude a role for the trans-sulfuration pathway in intracellular synthesis of cysteine given that the *Cystathionine Beta-Synthase* (*CBS*) mRNA was significantly upregulated only in this cell line upon FR054 treatment. Altogether these findings suggest that FR054 alone increases in both cell lines SLC7A11 glutamate/cystine exchange and intracellular GSSG level. Conversely, when combined with ERA, in MiaPaCa-2 cells might promote cell death through reducing glutamate/cystine exchange and GSH synthesis while in BxPC3 its proapoptotic effect, despite linked to a significant reduction of glutamate/cystine exchange, does not appear related to a GSH levels reduction.

### The combined effect between FR054 and erastin on cell survival is associated with oxidative imbalance causing lipid peroxidation

To evaluate the effect of intracellular GSH modulation and oxidative stress on PDAC cells survival under single and combined treatments with ERA and/or FR054, we used, as schematized in Figure S3C, two different approaches: 24h of cotreatment with *N*-acetyl-L-cysteine (NAC, 5mM), a cysteine prodrug, able to replenish intracellular GSH levels, or 24h of cotreatment with BSO (5mM) to block de novo synthesis of GSH. NAC cotreatment, as measured by trypan blue vital assay, led to decreased cell death in both cell lines and in all the conditions analyzed as compared to control cells (Figure 5A, B). Conversely, cotreatment with BSO led to increased cell death in cells treated with ERA alone or in combination with FR054 (Figure 5A, B). Of note, in FR054-treated samples, in which previous data indicated a significant enhancement of SLC7A11 activity tightly associated with a great accumulation of GSSG, suggesting a more functionally GSH biosynthetic pathway (Figure 2, 4), the effect of BSO was neglectable.

**Figure 5.**
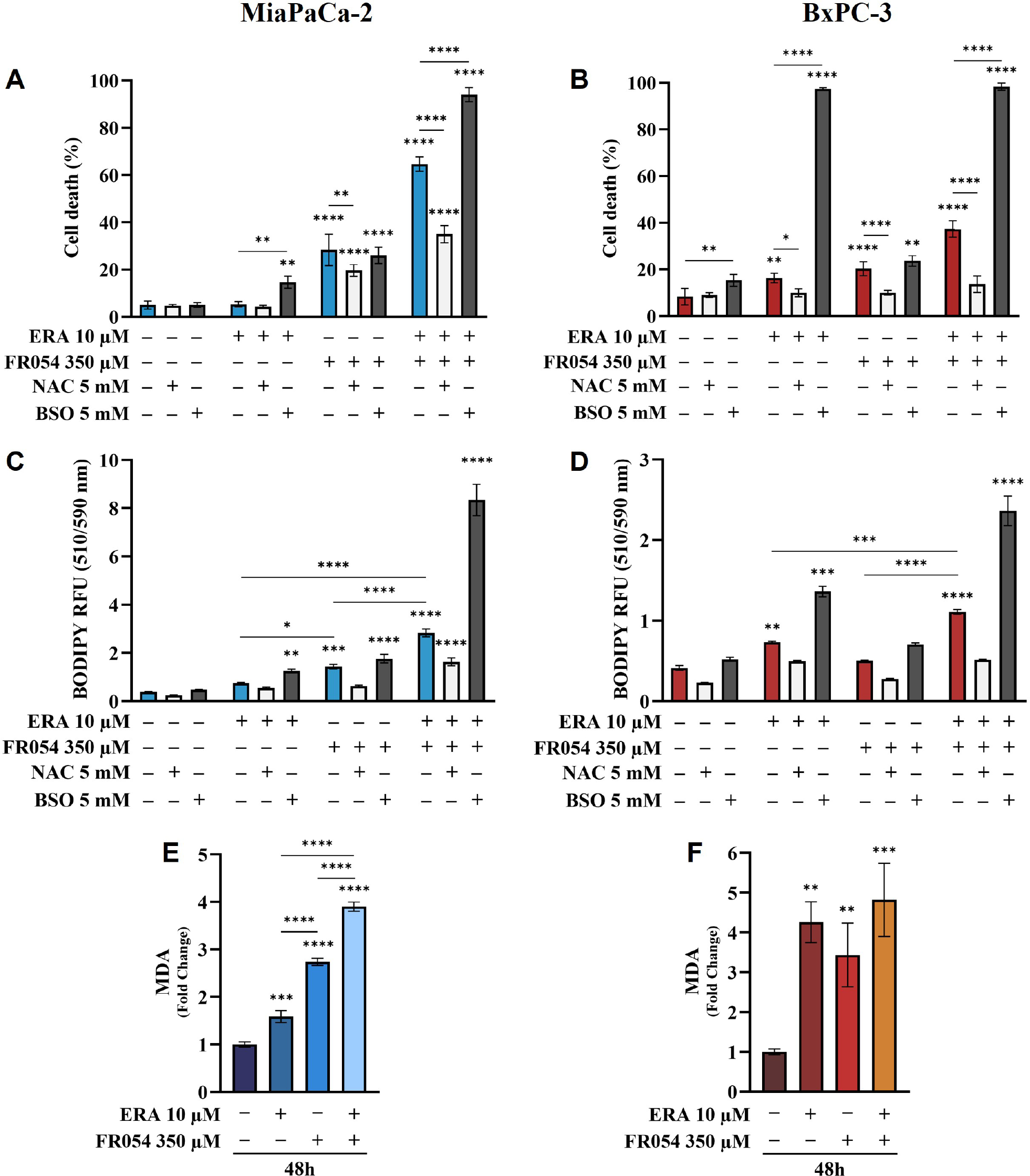
Variation of intracellular level of GSH by using NAC or BSO preserves or impairs respectively PDAC cell vitality by decreasing or increasing lipid peroxidation in PDAC cells. **(A)** and (B) Death cell percentage data (trypan blue assay) of MiaPaCa-2 and BxPC3 cells after the indicated treatments; (C) and (D) Intracellular Lipid-ROS levels in MiaPaCa-2 and BxPC3 cells after the indicated treatments; (E) and (F) malondialdehyde (MDA) assay to evaluate the lipid peroxidation in MiaPaCa-2 and BxPC3 cells after the indicated treatments; *p < 0.05, **p < 0.01 ***p < 0.001, ****p < 0.0001. The death cell percentage data and MDA assay data are presented as mean ± SD from three independent experiments, while intracellular Lipid-ROS data are presented as mean ± SEM from three independent experiments.

Based on our transcriptional data, revealing a role of ferroptosis in PDAC cells after FR054 treatment, given the significant effect of FR054 on NRF2 pathway activation as well as the noteworthy impact of BSO on PDAC cell survival when combined with ERA alone or in combination with FR054, we next sought to measure lipid peroxidation (Lipid-ROS) at 48h as a hallmark of cystine depletion as well as ferroptosis by using the fluorescent probe C11-BODIPY. As expected from previous results, treatment with ERA or FR054 alone increased fluorescence, especially in ERA-sensitive BxPC3 and slightly in FR054-treated MiaPaCa-2 cells. Conversely, combined treatment induced a significant increase in fluorescence in both cell lines (Figure 5C, D). As expected by cell survival data, NAC and BSO treatments reduced and increased Lipid-ROS, respectively, especially in cells treated with the drug combination, suggesting that these cells are more sensitive to intracellular redox state. To further demonstrate lipid peroxidation, we measured the malondialdehyde (MDA), an end-product of lipid peroxides, generated during ferroptosis and other oxidative stress context. As shown in Figure 5E and F, in both cell lines MDA levels, almost resembled C11-BODIPY staining, given that increased upon all treatments and especially in combined-treated samples. Thus, these findings indicate that FR054, increasing cell dependence for the SLC7A11-GSH axis, cause a significant increase in Lipid-ROS when combined with ERA and this effect occurs in both cell lines.

### Stable-isotope-assisted metabolomics analysis of central carbon metabolism indicates that the FR054 enhancing effect on ferroptosis is associated with a reduction of glucose entry into the TCA cycle and an increased glutaminolysis

To further investigate the underlying basis for the role of FR054 in increasing ERA sensitivity, we conducted a stable-isotope-assisted metabolomics analysis comparing all cell lines after the different treatments as described in Figure S3D. Metabolic analysis was performed by labeling the different cell lines with either [U-^13^C^6^]-glucose or [U-^13^C^5^]-glutamine and quantified isotopic enrichment in form of mass isotopomer distributions (MID). To analyze glycolysis, TCA cycle, and glutaminolysis, we analyzed the glucose or glutamine carbon contribution to the generation of pyruvate, citrate, *α*-ketoglutarate, and glutamate. In both cell lines, we observed a significant reduction of M4 and M5 citrate when cells were treated with FR054 alone or in combination with ERA, suggesting a reduced activity of both pyruvate dehydrogenase (PDH) and pyruvate carboxylase (PC) (Figure 6A, 7A, Figure S4A). We also observed this reduction in the M2 of *α*-ketoglutarate (Figure 6A, 7A), a key TCA cycle intermediate, pointing towards an overall of reduction of glucose entry into the TCA cycle. Indeed, in both cell lines, we observed a significant decrease in the glucose carbon contribution for their synthesis (Figure 6C, 7C). The effect of FR054, however, was much stronger in MiaPaCa-2 cells compared to BxPC3. Decreased glucose contribution suggests that cells were more dependent on glutamine for TCA cycle anaplerosis. In fact, analysis of the same metabolites in cells labeled with [U-^13^C^5^]-glutamine, indicated significantly higher enrichment of M5 glutamate, M5 *α*-ketoglutarate, and M4 citrate upon all treatments and especially in combined treatment (Figure 6B and 7B, Supplementary Figure S4B), suggesting that FR054 enhances glutaminolysis, as confirmed by the significant increase of relative glutamine carbon contributions for their synthesis (Figure 6C and 7C). Similar to the [U-^13^C^6^]-glucose results, the effect was much stronger in MiaPaCa-2 cells compared to BxPC3. An increase in labeled M3 pyruvate was observed only in ERA-treated samples in MiaPaCa-2 cells, suggesting only a slight alteration of malic enzyme activity in ERA-treated cells (Figure 6B). Nonetheless, analysis of the glutamine carbon contribution confirmed that in FR054 treated BxPC3 cells, glutamine become more relevant but to a lesser extent as compared to K-Ras mutated MiaPaCa-2 (Figure 7C). Altogether metabolomics analysis suggests that in both cell lines, treatment with FR054 alone or in combination, with a different magnitude, reduces glucose utilization through the TCA cycle. Here, we report that especially in MiaPaCa-2 cells, FR054 treatment significantly enhances cell dependence on glutamine and glutaminolysis, in line with previous findings describing glutaminolysis as a mechanism to increase ferroptosis cell death (41).

**Figure 6.**
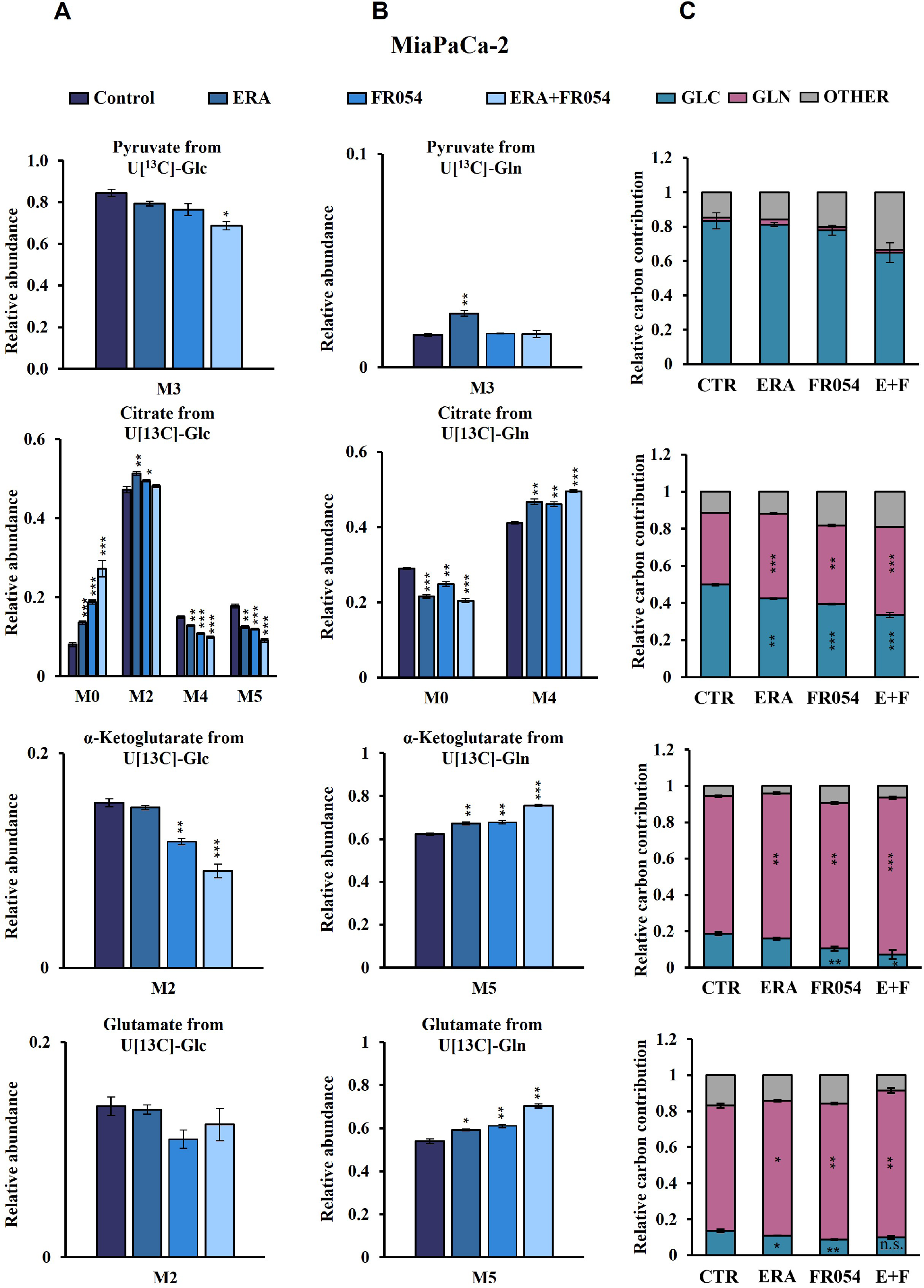
Combined treatment in MiaPaCa-2 cells favours glutaminolysis over glycolysis. **(A)** and (B) 13C labelling of pyruvate, citrate, a-ketoglutarate and glutamate from MiaPaCa-2 cells incubated in [U-^13^C^6^]-glucose or [U-^13^C^5^]-glutamine medium for 48h; (C) Relative carbon fractional contribution of glucose and glutamine to pyruvate, citrate, a-ketoglutarate and glutamate formation. *p < 0.05, **p < 0.01 ***p < 0.001. The data are presented as mean ± SEM from three independent experiments.

**Figure 7.**
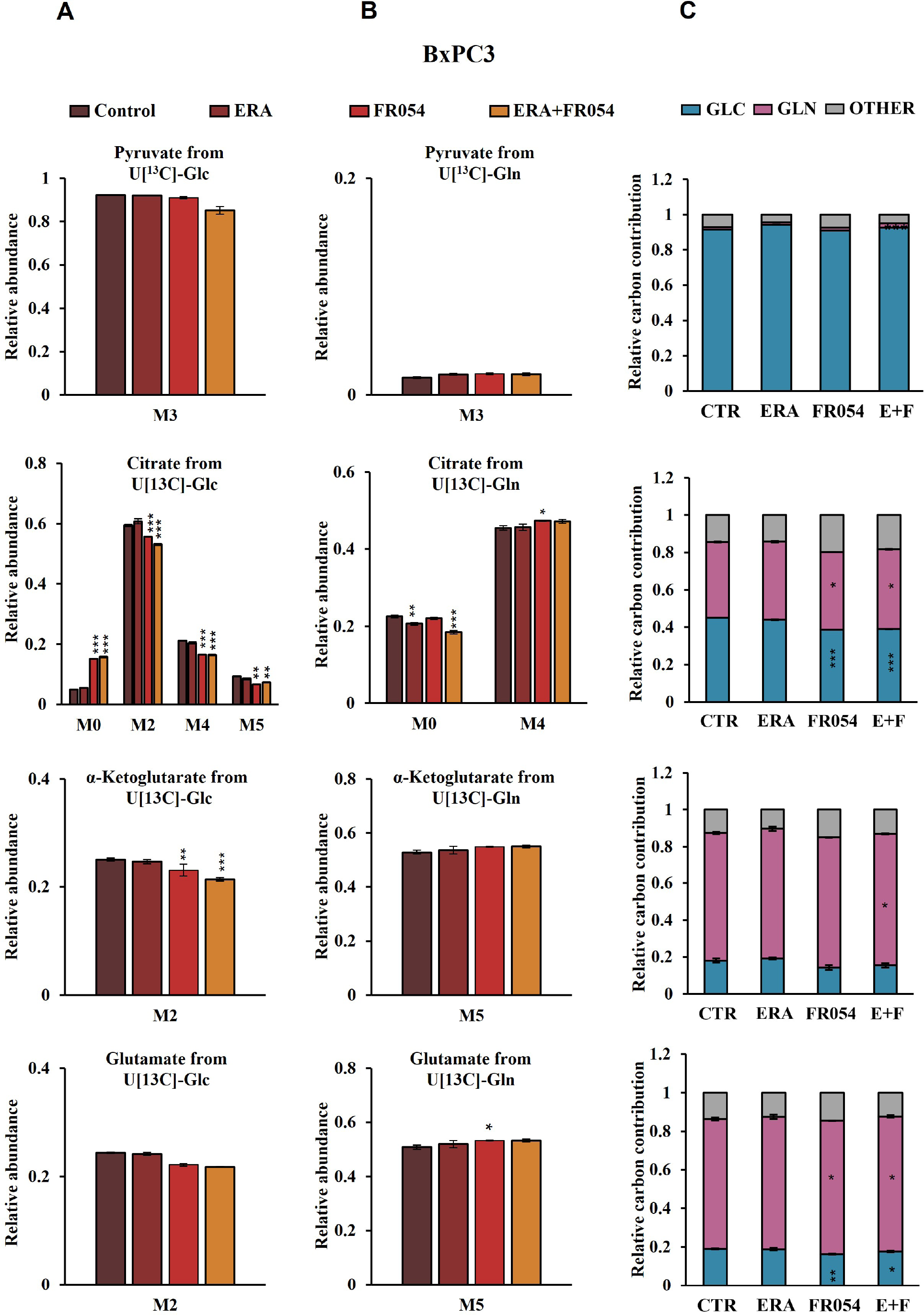
Combined treatment in BxPC3 cells reduces glucose utilization through the TCA cycle and slightly increases glutaminolysis. **(A)** and (B) 13C labelling of pyruvate, citrate, a-ketoglutarate and glutamate from BxPC3 cells incubated in [U-^13^C^6^]-glucose or [U-13C5]-glutamine medium for 48h; (C) Relative carbon fractional contribution of glucose and glutamine to pyruvate, citrate, a-ketoglutarate and glutamate formation. *p < 0.05, **p < 0.01 ***p < 0.001. The data are presented as mean ± SEM from three independent experiments.

### Combined treatment between ERA and FR054 induces enhancement of ferroptosis in PDAC cell lines independent on K-Ras mutation

To further reinforce the notion that ERA and FR054 treatment could cooperate in inducing ferroptosis in PDAC cell models, we treated human pancreatic cancer PANC1 cells, characterized by K-Ras mutation and a great sensitivity to cysteine depletion, with ERA and FR054 alone or in combination along a time course of 72h. We observed a significant effect of all treatments on proliferation (Figure 8A) and a time-dependent effect on cell death (Figure 8B). In both type of analysis, the combinatorial treatment induced the stronger outcome as confirmed also by morphological cell analysis, in which was observed a cell swelling before cell disruption (Figure 8C). Next, we analyzed glutamate and cystine levels, confirming that ERA alone or in combination significantly inhibited SLC7A11 function (Figure 8D, E) as well as FR054 alone induced a significant increase in SLC7A11 function. To evaluate the downstream effects of SLC7A11 modulation by the different treatments, we measured GSH and GSSG levels. The effect of all treatments significantly reduced GSH levels and such reduction was much stronger compared to MiaPaCa-2 and BxPC3 cells (Figure 8F). Of note, GSSG levels were significantly modulated by all treatments and to the same extent as in MiaPaCa-2 cells, further confirming the ability of ERA and FR054 to decrease or increase GSSG level, respectively (Figure 8G). Then we examined whether upon all treatments and adding NAC or BSO, PANC1 cells promoted lipid ROS and MDA generation. Single and combined treatments significantly increased C11-BODIPY fluorescence and intracellular MDA (Figure 8H, I). NAC and BSO treatment, as previously observed for the other two PDAC cell lines, significantly reduced or increased C11-BODIPY fluorescence, suggesting that also in PANC1 cells the combined treatment enhanced ferroptosis. Remarkable, our findings confirmed the great sensitivity of PANC1 cells to ERA treatment, as previously described, sensitivity that was further enhanced by FR054. Thus, the enhancer role of FR054 in ferroptosis could be attributed to the boosting of lipid peroxidation.

**Figure 8.**
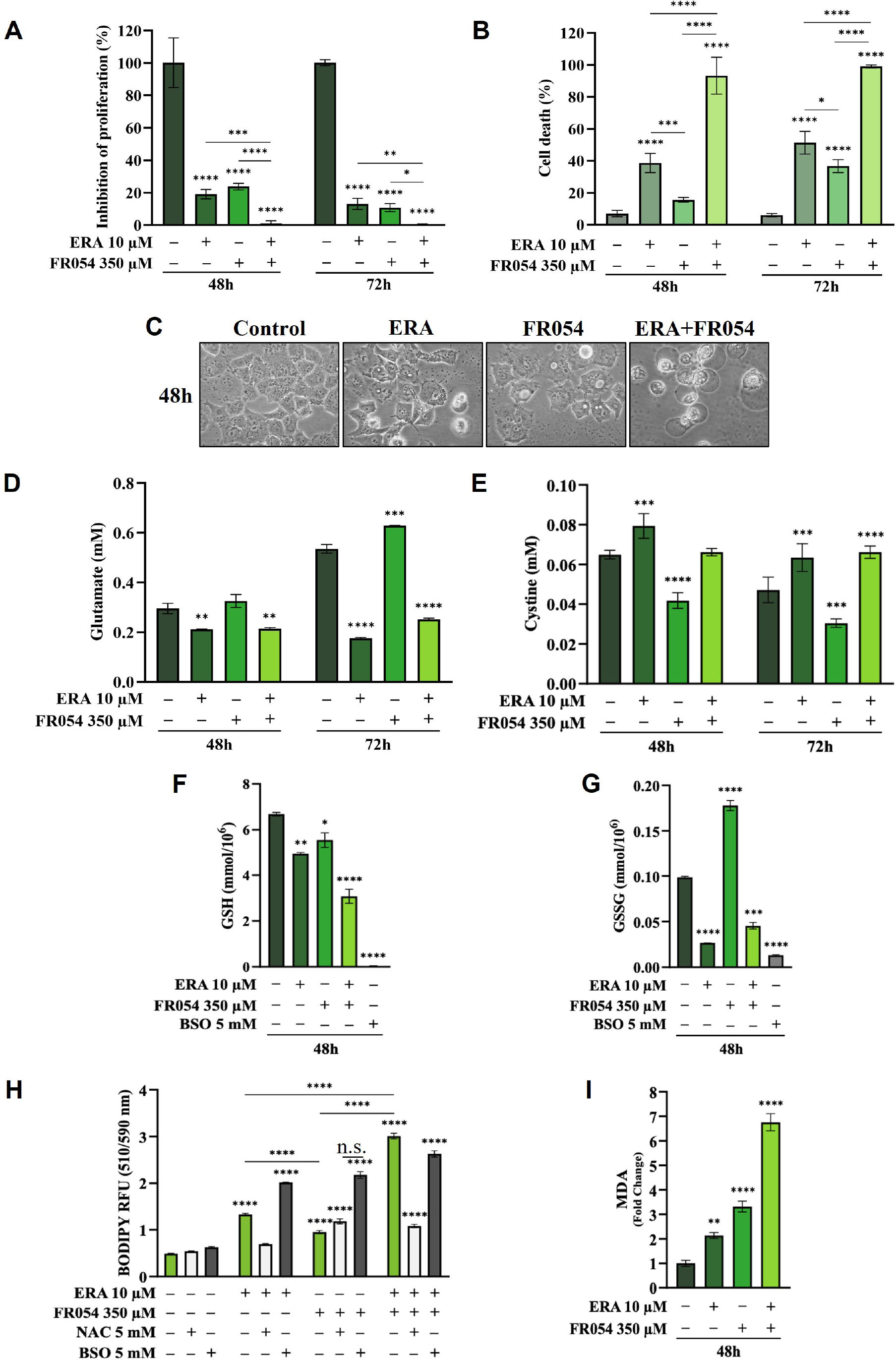
Combined treatment with FR054 and erastin enhances cell proliferation arrest and death of PANC1 cells by interfering with SLC7A11 activity, GSH metabolism and causing lipid peroxidation. **(A)** Trypan blue cell count in percentage after 48h and 72h of treatment with FR054, erastin (ERA) 24h or 48h, or their combination; (B) Cell death expressed as percentage (trypan blue assay) after 48 and 72h of treatment with FR054 alone or in combination with ERA; (C) Bright-field images of PANC1 cells treated as (A) taken 48h after the treatment; (D) and (E) HPLC evaluation of glutamate and cystine growth medium concentration, respectively, after the different treatments as indicated in figure; (F) GSH and (G) GSSG intracellular analysis upon the different treatments as indicated in figure; (H) intracellular Lipid-ROS levels in PANC1 cells after the indicated treatments; malondialdehyde (MDA) assay to evaluate the lipid peroxidation in PANC1 cells after the indicated treatments. *p < 0.05, **p < 0.01 ***p < 0.001, ****p < 0.0001. The data are presented as mean ± SD from three independent experiments. Only the data of GSH and GSSG concentration, and the data of Lipid-ROS levels are presented as mean ± SEM from three independent experiments.

To further dissect whether the mechanism by which FR054 increased ERA sensitivity in PANC1 cells as compared to other two PDAC cell lines, we conducted also in PANC1 a stable-isotope-assisted metabolomics analysis as previously described. In PANC1 cells, we observed a significant reduction of M4 and M5 citrate when cells were treated with ERA and FR054 alone or in combination, suggesting a reduced activity of both PDH and PC (Figure S7A). We also observed this reduction in the M2 of *α*-ketoglutarate and glutamate (Figure S7A), further suggesting an overall of reduction of glucose entry into the TCA cycle as demonstrated also by the significant decrease in the glucose carbon contribution for their synthesis (Figure S7C). Next, we analysed the same metabolites in PANC1 cells labelled with [U-^13^C^5^]-glutamine. A significant higher enrichment of M5 glutamate, M5 *α*-ketoglutarate, and M4 citrate upon all treatments and especially in combined treatment was observed (Figure S7B), suggesting that the combined treatment enhances glutaminolysis, as confirmed by the significant increase of relative glutamine carbon contributions for their synthesis (Figure S7C). Altogether, metabolomics analysis indicated that also in PANC1 cells, as observed in MiaPaCa-2 cells, the combined treatment reduces glucose utilization through the TCA cycle and significantly enhances cell dependence on glutamine and glutaminolysis, suggesting a role of glutaminolysis in ferroptosis induced by the combined treatment.

## Discussion

Previous studies suggested that PDAC exhibit extensive metabolic reprogramming necessary to support survival and growth and that such metabolic changes could represent new possible therapeutic targets (42). On the other hand, it has been recently shown that cysteine depletion induces PDAC cells death through a process named ferroptosis, a novel mechanism of non-apoptotic cell death, recently identified as another promising tumour therapeutic strategy (43). In this study, we investigated the anticancer effect of FR054, an inhibitor of the metabolic route named HBP, found altered in PDAC (12,13,44,45), in combination with ERA, a specific inhibitor of the antiporter SLC7A11, able to reduce cysteine uptake, and hence to stimulate ferroptosis. We show that such a combination, tested for the first time to the best of our knowledge, induces a significant enhancement of the ERA effect causing massive cell death, latter characterized by both apoptosis and ferroptosis. Of note, we show that the combination stimulates as compared to single treatments, an ER stress-dependent mechanism of cell protection, namely the ATF4-NRF2-SLC7A11-GSH axis, that causes an increased dependence on SLC7A11 activity, whose inhibition with ERA leads to massive cell death. Of note, although recent reports indicate both positive and negative association between ER stress and ferroptosis (46,47), the mechanism supporting a contribution of ER stress in ferroptosis remains to be further clarified.

Here, by using RNA-seq data and GSEA we identified that UPR activation, due to HBP inhibition, stimulates a wide transcriptional response involving several key regulators of cellular antioxidative response including the transcription factors NRF2, that lies at the center of a complex regulatory network governing cellular oxidative stress. Indeed, we identified several mRNAs involved in GSH synthesis, ROS detoxification, iron metabolism, and metabolic enzymes all upregulated especially in oncogenic K-Ras expressing MiaPaCa-2 cells as compared to wild-type expressing K-Ras BxPC3 cells. Such antioxidant response activated by UPR has been already described, indeed, accumulation of unfolded proteins causes an internal ER oxidative stress at which ER reacts by activation of a complex response among which is the induction of NRF2 (48). Importantly, cellular oxidative stress is able to activate ER stress as well (49). In addition, UPR modulates autophagy and ROS metabolism. More recent findings indicate that it may have an important impact also on cellular iron homeostasis i.e. by modulating the gene expression of ferroportin and ferritin (50). Therefore, in line with our previous findings, we can assume that FR054, interfering with protein folding, is a potent UPR inducer, as confirmed by our transcriptional data in which we observed upregulation of some UPR targets in both cell lines such as DDIT3 (CHOP), ATF4, ATF3 and CHAC1 (Figure 2). Intriguingly our transcriptional analysis showed that FR054 treatment caused a rather contrasting effect since we observed a significant upregulation of several mRNAs encoding for proteins involved in GSH metabolism among which SLC7A11 and HMOX1, latter an ER-resident protein, whose expression has been related to induction of ferroptosis (51,52).

Based on these findings and considering that ferroptosis can be exploited and then applied in clinical treatment strategies, we reasoned that a ferroptosis inducer, ERA, could significantly enhance the effect of FR054 in stimulating cell death. Indeed, in all the PDAC cell lines described in the manuscript, the combined treatment as compared to single ones caused an earlier cell proliferation arrest and then almost 100% of cell death. Of interest, analysis of the pro-apoptotic protein CHOP confirmed its expression in both ERA and FR054 treated cell samples, therefore substantiating a possible apoptotic mechanism activated by the single and combined treatments. On the other hand, another key factor involved in UPR response is ATF4. Previous studies have demonstrated that ATF4 has a protective role against ferroptosis or amino acid deprivation in cancer cells (26,30,53). However, other studies also indicated that ATF4 may contribute to cell death under the same types of stress (53–55). Therefore, ATF4 has a dual role in cell death and consequently, its final effect as a regulator of ferroptosis appears to be cell context-dependent.

Here we demonstrated that in ERA resistant-MiaPaCa-2 cells, ERA treatment was able to induce the UPR, as confirmed by increased GRP78 and CHOP expression, a pro-survival anti-oxidative response, as confirmed by NRF2 enhanced expression and above all a high level of ATF4, imputable to cystine deprivation. Conversely, FR054 in the same cells, also induced UPR and the anti-oxidative response but not a comparable level of ATF4, since no cystine depletion was observed upon this treatment. Strikingly, MiaPaCa-2 cells treated with the combination showed at an early time point (48h), a significant stronger activation of UPR and of the anti-oxidative response but a lower level of ATF4 as compared to ERA. However, at a later time point (72h), only the UPR remained significantly active as the anti-oxidative response as well as ATF4 significantly declined. On the other hand, the regulation of the same proteins in BxPC3 cells indicated their different behavior following the different treatments. Indeed, in ERA and FR054 treated cells the UPR, ATF4, and anti-oxidative responses were slowly activated and in a minor measure as compared to MiaPaCa-2 cells. Nonetheless, also in this cell line, the combined treatment induced a persistent UPR and a significant decrease of the anti-oxidative response as well as of ATF4, suggesting a minor cysteine depletion, as we observed, and slower intracellular oxidative stress. From these data, we can argue that the lower sensibility to ERA of MiaPaCa-2 cells as compared to BxPC3 cells, as previously described (24), must be probably ascribed to their ability to induce an ATF4-dependent response that together with NRF2 activation is able to avoid the cytotoxic effect of ERA. Instead, in BxPC3 cells, despite we observed only a significant reduction of extracellular glutamate and almost any change in extracellular cystine accumulation, ERA treatment induced almost 30% of cell death and a significantly reduced ATF4 accumulation, probably justifying their greater sensitivity. The ATF4 behavior in cells treated with the drug combination is quite interesting. Indeed, since ATF4 expression is associated with cancer cells resistance to ERA, in particular in MiaPaCa-2 cells, its decreased expression at later time point well fits with the highest levels of cell death observed (Figure 3E). Its reduction may be ascribed to the prolonged UPR activation, due to FR054 treatment under stressing conditions. Indeed, CHOP-dependent apoptosis is characterized by a specific restoration of protein translation, in order to transcribe pro-apoptotic genes (56,57). In a such condition, ATF4 becomes less active, due either to the expression of the inactive kinase *Tribbles Pseudokinase 3* (*TRIB3*), which acts as a negative feedback regulator of the ATF4-dependent transcription during the UPR (58), or is also degraded through a ubiquitination dependent mechanism (59,60). In this regard, we would underline that FR054 is able to induce transcriptionally *TRIB3* as results from our RNA-seq data (log^2^FoldChange: 9,53 and 1,71 in MiaPaCa-2 and BxPC3 cells, respectively) confirming the inactivation of ATF4’s function. Furthermore, it has been recently shown that Glycogen Synthase Kinase 3 Beta (GS3K) can target ATF4 for degradation. Such mechanism is controlled by the state of activation of AKT Serine/Threonine Kinase (AKT). Indeed, attenuation of the AKT pathway, leading to GSK3 activity, causes ATF4 degradation (61). Noteworthy, our previous findings demonstrated that FR054 can significantly reduce AKT activation in PDAC cells (12), therefore corroborating this mechanism as a possible way to induce degradation of ATF4 and to avoid its protective role. Our findings indicated also that FR054 reduces also NRF2 expression (Figures 4A, B). Interestingly, GSK3 activity controls also NRF2 ubiquitination and degradation suggesting a common mechanism of action on both NRF2 and ATF4 (62,63). Furthermore, analysis of KEAP1 expression, a key protein in NRF2 degradation, indicated that KEAP1, in both cell lines, is significantly induced in an FR054-dependent fashion, as shown in Figure 4A (compare lanes 3, 4, 7 and 8 with lanes 1, 2, 5 and 6) and Figure 4B (compare lanes 3, 4, 7 and 8 with lanes 1, 2, 5 and 6). The parallel increase of both NRF2 and KEAP1 suggests, as previously observed, a post-induction of KEAP1 in order to turn off the NRF2 signaling (64) that ultimately well fit with the increased cell death observed in FR054-treated samples at 72h. Regarding the observed parallel increase of NRF2 and KEAP1, it is worth of note to observe that some authors demonstrated that persistent accumulation of NRF2 is harmful (65,66). In this regard, some reports indicated that NRF2 is able to activate an auto-regulatory loop by inducing the expression of KEAP1 (67,68). Intriguingly, O-linked N-acetylglucosamine (GlcNAc) transferase (OGT) activation, the only enzyme deputy to protein O-glycosylation, has been associated with NRF2-dependent stress response, since OGT inhibition may facilitate NRF2 activation (69,70). The mechanism described for such an OGT-dependent NRF2 regulation is the O-glycosylation status of KEAP1 (71). Indeed, O-GlcNAcylation of KEAP1 is required for the efficient ubiquitination and degradation of NRF2 (71). Of interest, NRF2 degradation and ferroptosis have been recently linked through protein O-glycosylation. In particular, protein O-glycosylation regulates ferritinophagy and mitochondria behaviors to dictate ferroptosis, in particular it has been shown that inhibition of O-glycosylation promotes ferroptosis (72). In this scenario, we may hypothesize that ERA and FR054 cotreatment causes an overactivation of NRF2, following UPR and oxidative stress increase, as well as protein de-glycosylation (11,12), that eventually will activate a KEAP1-dependent regulatory negative loop that causes NRF2 degradation and cellular oxidative stress. This hypothesis is supported by our observation of a robust activation of NRF2 and KEAP1 in combined treatment in both cell lines (Figure 4A, 4B, Figure S6, S7), an increased lipid peroxidation (Figure 5C, D, E, F) and cell death (Figure 5A, 5B). Despite we do not test yet protein expression in PANC1 cells, our hypothesis is strongly supported also in this cell line as shown in Figure 8 in which the combined treatment induces a significant increase in cell death, inhibits SLC7A11 activity, and increases ferroptosis markers.

In cancer cells, inhibition of system SLC7A11-mediated cystine uptake by ERA, may be sufficient to initiate ferroptosis by interfering with GSH and GSSG intracellular levels. Our results in the three PDAC cell lines indicated that ERA was able to induce a small but significant decrease in GSH level only in PANC1 and BxPC3 cells (Figure 4I, J and Figure 8F, G), given that in MiaPaCa-2 cells the decrease was unimportant. However, these findings agree with previous research, showing that PANC1 and BxPC3 cells are more sensitive to imidazole ketone erastin (IKE), an erastin analogue, as compared to MiaPaCa-2 cells, suggesting that the cell ability to avoid ERA effect is associated to GSH intracellular levels (73). Accordingly, combined treatment, able to significantly increase a ferroptosis mechanism of cell death (Figure 5C, D, E, F and 8H and J), significantly reduced GSH levels in MiaPaCa-2 and PANC1 cells. Regarding the GSH level in BxPC3, must be underlined that although in combined treatment we observed a significant reduction and accumulation of extracellular glutamate and cystine, respectively, we did not observe a GSH reduction, suggesting a more complex mechanism causing ferroptosis in this cell line. Worth of note, in all three cell lines, upon ERA and combined treatments, we observed a consistent and significant decrease of GSSG. This result is both in agreement (73–76) and in disagreement with other published observations because other researchers have shown that GSH depletion is accompanied by high GSSG levels. While we do not have an explanation about our different results, we would underline that in our experimental conditions also BSO treatment caused a significant reduction of both GSH and GSSG as compared to control cells, suggesting that the decline of total GSH may have a role in ferroptosis onset (76,77). Whether the GSSG level plays a role in the execution of ferroptosis remains unclear. An almost completely opposite behavior was observed in FR054-treated cells. Indeed, in line with the transcriptional data and the glutamate/cystine measurements, the effect of FR054 on GSH level as compared to control cells varied among cell lines (no change in MiaPaCa-2 cells, slightly high in BxPC3 cells and slightly low in PANC1 cells). Nevertheless, in all three cell lines upon FR054 treatment, the GSSG levels significantly increased suggesting that these cells were actively using GSH for cellular detoxification maybe to restore protein folding and cope with UPR stress (78). In this regard, a recent manuscript has been proposed an important role for GSSG in controlling Endoplasmic Reticulum (ER) function. Indeed, it has been shown that increased GSSG level may, directly or indirectly, alters ER oxidative protein maturation and protects ER homeostasis (79). How the level of GSSG may control ER homeostasis and ferroptosis are not the topic of this manuscript, but since few authors addressed this role, it could be the topic for future investigation.

Nevertheless, the cell death mechanism observed in all the PDAC cell lines and in particular in combined treatment is clearly dependent on GSH levels, since we observed that cell survival was almost completely restored or significantly reduced by adding NAC or BSO, respectively (Figure 5A, B). Moreover, analysis of lipid peroxidation and MDA confirmed the induction of a ferroptosis mechanism of cell death especially in combined treatment (Figure 5C, D, E, F and Figure 8H, I). Noteworthy, lipid peroxidation was observed also in FR054 treated cells, which was consistent with previous findings showing a crosstalk between the two processes, ER stress and ferroptosis, in controlling cell death through the activation of the PERK-eIF2α-ATF4-CHOP axis (80) or PERK-Nrf2-HMOX1 axis (46). It was intriguing that HMOX1 was identified as one of the most regulated mRNA in both cell lines upon FR054 treatment (Figure 1 and Figure 2), inferring that accumulation of ferrous iron (Fe^2+^), generated by HMOX1 activity, could cause further lipid peroxidation in combined treatment. Indeed, western blot analysis indicated that in both MiaPaCa-2 and BxPC3 cells was also induced at protein level (Figure 4A). Worth of note, we have to highlight that although in both cell lines, it was significantly induced at mRNA level upon FR054 treatment, at protein level was detected only in MiaPaCa-2 cells. We do not have an explanation for this discrepancy, but we would emphasize that all the other mRNAs tested by western blot, specifically ATF4, DDIT3/CHOP, NRF2, SLC7A11 were modulated similarly at both mRNA and protein level.

The role of cellular metabolism in ferroptosis execution has been addressed in several studies as reviewed in (81) and in fact, an increasing number of metabolic pathways appear to converge on ferroptosis. Intriguingly, we show that in all the PDAC cell lines the combined treatment causes a metabolic rewiring leading to a reduction of glucose entry into the TCA cycle. This effect, but to a lesser extent, was observed also following the single treatments (Figure 6A, 7A and Figure S7A). The effect well correlates with our findings in which we observed that FR054 treatment upregulates several genes of the glycolytic branches able to generate NADPH such as *Glucose-6-Phosphate Dehydrogenase* (*G6PD*) and *6-Phosphogluconate Dehydrogenase* (*PGD*) of the Pentose Phosphate Pathway, *Phosphoserine Aminotransferase 1 (PSAT1)* and *Phosphoglycerate Dehydrogenase (PHGDH)* of the serine biosynthetic pathway, necessary for the recycling of the oxidized GSH through GSR activity in order to maintain the cellular redox homeostasis (82). Furthermore, we show that such glycolytic rewiring is associated with a concurrently increase in glutamine dependence of PDAC cells especially in K-Ras mutated MiaPaCa-2 and PANC1. The role of glutamine in ferroptosis is quite complex, however, some authors showed that glutamine drives ferroptosis in combination with cystine deprivation through glutaminolysis. Indeed, it has been proposed that inhibition of SLC7A11 activity, increasing glutaminolysis, causes a greater glutamate conversion into *α*-KG that boosts the TCA cycle, the mitochondria activity, the fatty acid synthesis, and the generation of ROS that eventually altogether concur to ferroptosis (83,84). Importantly, glutaminolysis alone is not able to trigger ferroptosis pointing out that ferroptosis may happen only when glutaminolysis is activated in a cysteine depletion condition (84–86). In contrast, it has been also shown that ferroptosis is further accelerated by inhibition of ETC complexes and OXPHOS (75,87). The discrepancy between previous studies and ours can be explained by the differences in cell type, ferroptosis inducers, and exposure time. Particularly, activation of ferroptotic signaling in response to ferroptosis inducers (RSL3, erastin, and cysteine-deprivation) might be mediated through different mechanistic pathways in mitochondria. On the other hand, other authors have shown that ferroptosis sensitivity is associated also with metabolic compartmentalization. Indeed, genetic silencing and pharmacological inhibition of glutaminolysis through GLS2, but not GLS1, or GOT1 have been shown to inhibit erastin-induced ferroptosis (88–90). Furthermore, also in our study, meanwhile the combined treatment was able to increase in all PDAC cells ferroptosis, we observed some differences between K-Ras mutated cell lines and K-Ras wild-type cells, especially about glutaminolysis activation (compare Figure 6, Figure 7 and Figure S7). Therefore, the differences emerging from our and other studies possible reflect the incomplete understanding of how various factors dictate sensitivity to ferroptosis.

In conclusion, our findings indicate that FR054 alone or in combination with ERA is able to modulate the oxidative stress response and central carbon metabolism, favoring glutaminolysis over glycolysis, that in a condition of SLC7A11 inhibition following ERA treatment, causes unbalance in cellular redox state, GSH depletion, lipid-Ros accumulation and finally ferroptosis. Based on our in vitro data we propose FR054 together with a drug able to modulate the intracellular level of cysteine, such ERA, may become a novel candidate for in vivo ferroptosis induction for the targeted killing of PDAC cells.

### Limitations

The scope of our study was to identify possible pathways or proteins whose inhibition/activation could synergize with FR054, a novel HBP inhibitor, in inducing PDAC cell death. However, our study contains certain limitations. First, a Q-PCR validation of some DEGs was not conducted in order to determine if some specific pathways were more activated by FR054 treatment. Second, we did not test the effect of a specific ferroptosis inhibitor such as Ferrostatin-1 to confirm the ability of the combined treatment in inducing ferroptosis. Third, we did not test our combination in immortalized pancreatic cells such as human pancreatic duct epithelial cells to support the use of the combination in preclinical in vivo models. Finally, fourth, we did not directly assess the mitochondrial activity and the role of metabolism compartmentalization, as we have done previously (91), in our cell models upon the different treatments, causing ferroptosis.

## 5 Author Contributions

Conception and design: AW and FC. Development of methodology: BZ, MG, TL, AP, LB, AW and FC. Performing of research: BZ, MB, TL, VB, AP. Analysis and interpretation of data: BZ, MG, TL, AP, LB, AW and FC. Writing the draft of the manuscript: FC. Review of the manuscript: BZ, VB, AW and FC. Revision of the manuscript: all authors. All authors contributed to the article and approved the submitted version.

## 9 Supplementary Material

Supplementary Table 1. DEGs identified in MiaPaCa-2 and BxPC3 cells upon FR054 treatment as compared to untreated cells.

Supplementary Figure S1. Table listing all significative enriched gene sets in treated MIAPaCa-2 cells. Supplementary Figure S2. Table listing all significative enriched gene sets in treated BxPC3 cells. Supplementary Figure S3. Experimental setting of the different treatments described in the main text. Supplementary Figure S4. The combined treatment of MiaPaca-2 cells with FR054 and erastin affects UPR and oxidative stress response proteins expression.

Supplementary Figure S5. The combined treatment of BxPC3 cells with FR054 and erastin affects UPR and oxidative stress response proteins expression.

Supplementary Figure S6. Isotopomers labeling for metabolomics study in MiaPaCa-2, BxPC3 and PANC1 cell lines.

Supplementary Figure S7. Combined treatment in PANC1 cells favors glutaminolysis over glycolysis

## 10 Data Availability Statement

The RNA-seq data have been deposited at NCB-GEO database, accession number GSE223303 (https://www.ncbi.nlm.nih.gov/geo/query/acc.cgi?acc=GSE223303).

