## Supplementary Figure 1 for "PGM3 inhibition Shows cooperative Effects With Erastin inducing Pancreatic cancer cell death via activation of the Unfolded Protein Response"

| # | pathway | NES | FDR | count |
|---|---|---|---|---|
| 1 | PHOTODYNAMIC THERAPY INDUCED UNFOLDED PROTEIN RESPONSE | 2.48E+00 | 0.00E+00 | 23 |
| 2 | CYTOPLASMIC RIBOSOMAL PROTEINS | 2.31E+00 | 6.57E-04 | 86 |
| 3 | PHOTODYNAMIC THERAPY INDUCED NFE2L2 NRF2 SURVIVAL SIGNALING | 2.13E+00 | 9.16E-03 | 23 |
| 4 | NRF2 PATHWAY | 2.11E+00 | 1.02E-02 | 104 |
| 5 | TRANSCRIPTIONAL ACTIVATION BY NRF2 IN RESPONSE TO PHYTOCHEMICALS | 2.03E+00 | 1.72E-02 | 13 |
| 6 | EXERCISEINDUCED CIRCADIAN REGULATION | 2.01E+00 | 1.92E-02 | 41 |
| 7 | UNFOLDED PROTEIN RESPONSE | 2.03E+00 | 2.04E-02 | 24 |
| 8 | SEROTONIN AND ANXIETYRELATED EVENTS | 1.98E+00 | 2.49E-02 | 6 |
| 9 | OXIDATIVE STRESS RESPONSE | 1.94E+00 | 3.45E-02 | 28 |
| 10 | NRF2ARE REGULATION | 1.93E+00 | 3.55E-02 | 21 |
| 11 | OREXIN RECEPTOR PATHWAY | 1.91E+00 | 4.01E-02 | 103 |
| 12 | MRNA PROTEIN AND METABOLITE INDUCATION PATHWAY BY CYCLOSPORIN A | 1.89E+00 | 4.86E-02 | 7 |
| 13 | VITAMIN DSENSITIVE CALCIUM SIGNALING IN DEPRESSION | 1.87E+00 | 5.61E-02 | 26 |
| 14 | PREIMPLANTATION EMBRYO | 1.83E+00 | 7.64E-02 | 37 |
| 15 | FERROPTOSIS | 1.82E+00 | 8.09E-02 | 59 |
| 16 | GENES RELATED TO PRIMARY CILIUM DEVELOPMENT BASED ON CRISPR | -1.86E+00 | 8.10E-02 | 89 |
| 17 | DEREGULATION OF RAB AND RAB EFFECTOR GENES IN BLADDER CANCER | -1.88E+00 | 8.90E-02 | 15 |
| 18 | BENZOAPYRENE METABOLISM | 1.80E+00 | 9.25E-02 | 7 |
| 19 | MIR517 RELATIONSHIP WITH ARCN1 AND USP1 | 1.77E+00 | 1.06E-01 | 5 |
| 20 | MRNA PROCESSING | 1.76E+00 | 1.15E-01 | 124 |
| 21 | TYPE I INTERFERON INDUCTION AND SIGNALING DURING SARSCOV2 INFECTION | -1.89E+00 | 1.27E-01 | 27 |
| 22 | WHITE FAT CELL DIFFERENTIATION | 1.73E+00 | 1.34E-01 | 29 |
| 23 | DRUG INDUCTION OF BILE ACID PATHWAY | 1.73E+00 | 1.37E-01 | 6 |
| 24 | EUKARYOTIC TRANSCRIPTION INITIATION | 1.72E+00 | 1.45E-01 | 40 |
| 25 | IRON METABOLISM IN PLACENTA | 1.70E+00 | 1.54E-01 | 10 |
| 26 | PROTEASOME DEGRADATION | 1.69E+00 | 1.58E-01 | 55 |
| 27 | ANTIVIRAL AND ANTIINFLAMMATORY EFFECTS OF NRF2 ON SARSCOV2 PATHWAY | 1.70E+00 | 1.60E-01 | 25 |
| 28 | HYPERTROPHY MODEL | 1.70E+00 | 1.64E-01 | 16 |
| 29 | TRANSCRIPTIONAL CASCADE REGULATING ADIPOGENESIS | 1.67E+00 | 1.68E-01 | 13 |
| 30 | COHESIN COMPLEX CORNELIA DE LANGE SYNDROME | 1.67E+00 | 1.70E-01 | 34 |
| 31 | NEPHROTIC SYNDROME | -1.91E+00 | 1.75E-01 | 38 |
| 32 | BLADDER CANCER | 1.67E+00 | 1.76E-01 | 36 |
| 33 | SEROTONIN AND ANXIETY | 1.65E+00 | 1.81E-01 | 9 |
| 34 | MIRNAS INVOLVED IN DNA DAMAGE RESPONSE | 1.63E+00 | 1.94E-01 | 21 |
| 35 | PARKINUBIQUITIN PROTEASOMAL SYSTEM PATHWAY | 1.62E+00 | 1.97E-01 | 60 |
| 36 | P53 TRANSCRIPTIONAL GENE NETWORK | 1.63E+00 | 1.98E-01 | 62 |
| 37 | GLUTATHIONE METABOLISM | 1.63E+00 | 1.99E-01 | 15 |
| 38 | CHROMOSOMAL AND MICROSATELLITE INSTABILITY IN COLORECTAL CANCER | 1.64E+00 | 2.01E-01 | 70 |
| 39 | HEMATOPOIETIC STEM CELL DIFFERENTIATION | 1.61E+00 | 2.01E-01 | 42 |
| 40 | NUCLEAR RECEPTORS METAPATHWAY | 1.63E+00 | 2.03E-01 | 234 |
| 41 | GANGLIO SPHINGOLIPID METABOLISM | 1.58E+00 | 2.41E-01 | 9 |
| 42 | STRIATED MUSCLE CONTRACTION PATHWAY | 1.58E+00 | 2.44E-01 | 24 |
| 43 | LET7 INHIBITION OF ES CELL REPROGRAMMING | 1.58E+00 | 2.50E-01 | 8 |

**Supplementary Figure S1. Table listing all significative enriched gene sets in treated MIAPaCa-2 cells.** Rankings based on FDR score. ± NES indicates upregulation or downregulation respectively of gene set in treated MIAPaCa-2 cells.

**Supplementary Figure S1**
