## Supplementary Figure 2 for "PGM3 inhibition Shows cooperative Effects With Erastin inducing Pancreatic cancer cell death via activation of the Unfolded Protein Response"

| # | pathway | enrichment | pvalue | count |
|---|---|---|---|---|
| 1 | FERROPTOSIS | 2.53E+00 | 0.00E+00 | 60 |
| 2 | OVERVIEW OF PROINFLAMMATORY AND PROFIBROTIC MEDIATORS | -1.99E+00 | 0.00E+00 | 66 |
| 3 | NETWORK MAP OF SARSCOV2 SIGNALING PATHWAY | -2.06E+00 | 0.00E+00 | 166 |
| 4 | SARSCOV2 INNATE IMMUNITY EVASION AND CELLSPECIFIC IMMUNE RESPONSE | -2.13E+00 | 0.00E+00 | 55 |
| 5 | ALLOGRAFT REJECTION | -2.26E+00 | 0.00E+00 | 48 |
| 6 | PHOTODYNAMIC THERAPYINDUCED NFE2L2 NRF2 SURVIVAL SIGNALING | 2.30E+00 | 1.00E-03 | 23 |
| 7 | TYPE II INTERFERON SIGNALING IFNG | -1.98E+00 | 1.00E-03 | 29 |
| 8 | CYTOKINES AND INFLAMMATORY RESPONSE | -1.91E+00 | 3.00E-03 | 17 |
| 9 | IMMUNE RESPONSE TO TUBERCULOSIS | -1.92E+00 | 3.00E-03 | 23 |
| 10 | TRANSCRIPTIONAL ACTIVATION BY NRF2 IN RESPONSE TO PHYTOCHEMICALS | 2.13E+00 | 4.00E-03 | 14 |
| 11 | NRF2ARE REGULATION | 2.10E+00 | 4.00E-03 | 22 |
| 12 | BENZOAPYRENE METABOLISM | 2.10E+00 | 4.00E-03 | 8 |
| 13 | ESTROGEN RECEPTOR PATHWAY | 2.07E+00 | 5.00E-03 | 11 |
| 14 | HIPPOMERLIN SIGNALING DYSREGULATION | -1.85E+00 | 9.00E-03 | 94 |
| 15 | SPINAL CORD INJURY | -1.86E+00 | 9.00E-03 | 89 |
| 16 | MIRNAS INVOLVED IN DNA DAMAGE RESPONSE | 2.02E+00 | 1.00E-02 | 23 |
| 17 | CHEMOKINE SIGNALING PATHWAY | -1.81E+00 | 1.50E-02 | 128 |
| 18 | MBDNF AND PROBDNF REGULATION OF GABA NEUROTRANSMISSION | -1.82E+00 | 1.50E-02 | 28 |
| 19 | INFLAMMATORY RESPONSE PATHWAY | -1.80E+00 | 1.60E-02 | 21 |
| 20 | CCL18 SIGNALING PATHWAY | -1.81E+00 | 1.60E-02 | 39 |
| 21 | AUTOPHAGY | 1.95E+00 | 2.00E-02 | 29 |
| 22 | MIR5093P ALTERATION OF YAP1ECM AXIS | -1.78E+00 | 2.00E-02 | 14 |
| 23 | DEVELOPMENT OF URETERIC COLLECTION SYSTEM | -1.77E+00 | 2.40E-02 | 50 |
| 24 | OXIDATIVE STRESS RESPONSE | 1.91E+00 | 3.00E-02 | 29 |
| 25 | NEURODEGENERATION WITH BRAIN IRON ACCUMULATION NBIA SUBTYPES PATHWAY | 1.90E+00 | 3.10E-02 | 44 |
| 26 | NICOTINE EFFECT ON DOPAMINERGIC NEURONS | -1.72E+00 | 4.00E-02 | 14 |
| 27 | COMPLEMENT SYSTEM | -1.73E+00 | 4.00E-02 | 58 |
| 28 | COVID19 ADVERSE OUTCOME PATHWAY | -1.73E+00 | 4.00E-02 | 9 |
| 29 | TYPE I INTERFERON INDUCTION AND SIGNALING DURING SARSCOV2 INFECTION | -1.74E+00 | 4.00E-02 | 26 |
| 30 | OLIGODENDROCYTE SPECIFICATION AND DIFFERENTIATION LEADING TO MYELIN COMPONENTS FOR CNS | -1.72E+00 | 4.20E-02 | 18 |
| 31 | PATHOGENESIS OF SARSCOV2 MEDIATED BY NSP9NSP10 COMPLEX | -1.73E+00 | 4.20E-02 | 14 |
| 32 | COMPLEMENT AND COAGULATION CASCADES | -1.71E+00 | 4.30E-02 | 40 |
| 33 | COMPLEMENT SYSTEM IN NEURONAL DEVELOPMENT AND PLASTICITY | -1.71E+00 | 4.40E-02 | 79 |
| 34 | SELECTIVE EXPRESSION OF CHEMOKINE RECEPTORS DURING TCELL POLARIZATION | -1.70E+00 | 5.10E-02 | 15 |
| 35 | PRIMARY FOCAL SEGMENTAL GLOMERULOSCLEROSIS FSGS | -1.69E+00 | 5.60E-02 | 65 |
| 36 | EBOLA VIRUS INFECTION IN HOST | -1.68E+00 | 6.10E-02 | 108 |
| 37 | GLUTATHIONE METABOLISM | 1.82E+00 | 6.20E-02 | 16 |
| 38 | GPCRS CLASS A RHODOPSINLIKE | -1.66E+00 | 6.80E-02 | 84 |
| 39 | NONGENOMIC ACTIONS OF 125 DIHYDROXYVITAMIN D3 | -1.66E+00 | 6.80E-02 | 62 |
| 40 | BURN WOUND HEALING | -1.66E+00 | 7.00E-02 | 82 |
| 41 | HIPPO SIGNALING REGULATION PATHWAYS | -1.65E+00 | 7.30E-02 | 79 |
| 42 | NEOVASCULARISATION PROCESSES | -1.65E+00 | 7.40E-02 | 34 |
| 43 | PLATELETMEDIATED INTERACTIONS WITH VASCULAR AND CIRCULATING CELLS | -1.64E+00 | 8.30E-02 | 12 |
| 44 | WNT SIGNALING | -1.63E+00 | 8.40E-02 | 96 |
| 45 | MIRNA TARGETS IN ECM AND MEMBRANE RECEPTORS | -1.63E+00 | 8.50E-02 | 24 |
| 46 | CARDIAC PROGENITOR DIFFERENTIATION | -1.63E+00 | 8.50E-02 | 34 |
| 47 | FOCAL ADHESION | -1.63E+00 | 8.60E-02 | 170 |

**Supplementary Figure S2**

| # | pathway | enrichment | pvalue | count |
|---|---|---|---|---|
| 48 | LUNG FIBROSIS | -1.62E+00 | 9.10E-02 | 40 |
| 49 | AIRWAY SMOOTH MUSCLE CELL CONTRACTION | -1.62E+00 | 9.20E-02 | 15 |
| 50 | NRF2 PATHWAY | 1.78E+00 | 9.30E-02 | 110 |
| 51 | ARYL HYDROCARBON RECEPTOR PATHWAY WP2873 | 1.77E+00 | 9.60E-02 | 38 |
| 52 | EDA SIGNALING IN HAIR FOLLICLE DEVELOPMENT | -1.61E+00 | 1.01E-01 | 12 |
| 53 | HAIR FOLLICLE DEVELOPMENT CYTODIFFERENTIATION PART 3 OF 3 | -1.60E+00 | 1.01E-01 | 64 |
| 54 | NEPHROTIC SYNDROME | -1.60E+00 | 1.02E-01 | 36 |
| 55 | BMP2WNT4FOXO1 PATHWAY IN PRIMARY ENDOMETRIAL STROMAL CELL DIFFERENTIATION | -1.60E+00 | 1.03E-01 | 11 |
| 56 | SARS CORONAVIRUS AND INNATE IMMUNITY | -1.59E+00 | 1.18E-01 | 17 |
| 57 | VITAMIN B12 METABOLISM | -1.58E+00 | 1.20E-01 | 37 |
| 58 | HEDGEHOG SIGNALING PATHWAY WP4249 | -1.58E+00 | 1.23E-01 | 39 |
| 59 | ESTROGEN METABOLISM | 1.72E+00 | 1.32E-01 | 14 |
| 60 | PROSTAGLANDIN SIGNALING | -1.56E+00 | 1.36E-01 | 24 |
| 61 | GENES TARGETED BY MIRNAS IN ADIPOCYTES | -1.57E+00 | 1.37E-01 | 10 |
| 62 | TGFBETA SIGNALING IN THYROID CELLS FOR EPITHELIALMESENCHYMAL TRANSITION | -1.56E+00 | 1.39E-01 | 16 |
| 63 | LNCRNA IN CANONICAL WNT SIGNALING AND COLORECTAL CANCER | -1.56E+00 | 1.39E-01 | 81 |
| 64 | PLURIPOTENT STEM CELL DIFFERENTIATION PATHWAY | -1.56E+00 | 1.42E-01 | 34 |
| 65 | PATHWAYS OF NUCLEIC ACID METABOLISM AND INNATE IMMUNE SENSING | -1.55E+00 | 1.42E-01 | 12 |
| 66 | ARRHYTHMOGENIC RIGHT VENTRICULAR CARDIOMYOPATHY | -1.55E+00 | 1.43E-01 | 54 |
| 67 | MAMMALIAN DISORDER OF SEXUAL DEVELOPMENT | -1.55E+00 | 1.48E-01 | 17 |
| 68 | VITAMIN B12 DISORDERS | -1.54E+00 | 1.58E-01 | 11 |
| 69 | NEURAL CREST CELL MIGRATION IN CANCER | -1.54E+00 | 1.59E-01 | 36 |
| 70 | HOSTPATHOGEN INTERACTION OF HUMAN CORONAVIRUSES INTERFERON INDUCTION | -1.53E+00 | 1.61E-01 | 32 |
| 71 | IL1 AND MEGAKARYOCYTES IN OBESITY | -1.53E+00 | 1.65E-01 | 20 |
| 72 | WNT SIGNALING IN KIDNEY DISEASE | -1.52E+00 | 1.80E-01 | 31 |
| 73 | GENES ASSOCIATED WITH THE DEVELOPMENT OF RHEUMATOID ARTHRITIS | -1.52E+00 | 1.81E-01 | 11 |
| 74 | PHOTODYNAMIC THERAPYINDUCED UNFOLDED PROTEIN RESPONSE | 1.67E+00 | 1.89E-01 | 25 |
| 75 | NCRNAS INVOLVED IN WNT SIGNALING IN HEPATOCELLULAR CARCINOMA | -1.50E+00 | 2.07E-01 | 74 |
| 76 | CELLS AND MOLECULES INVOLVED IN LOCAL ACUTE INFLAMMATORY RESPONSE | -1.50E+00 | 2.08E-01 | 12 |
| 77 | PI3K AKT mTOR SIGNALING PATHWAY AND THERAPEUTIC OPPORTUNITIES | 1.64E+00 | 2.12E-01 | 29 |
| 78 | ZINC HOMEOSTASIS | -1.50E+00 | 2.12E-01 | 29 |
| 79 | FIBRIN COMPLEMENT RECEPTOR 3 SIGNALING PATHWAY | -1.50E+00 | 2.14E-01 | 33 |
| 80 | ANGIOTENSIN II RECEPTOR TYPE 1 PATHWAY | -1.49E+00 | 2.24E-01 | 23 |
| 81 | MALIGNANT PLEURAL MESOTHELIOMA | -1.48E+00 | 2.28E-01 | 371 |
| 82 | TRANSCRIPTION COFACTORS SKI AND SKIL PROTEIN PARTNERS | 1.60E+00 | 2.32E-01 | 18 |
| 83 | AFLATOXIN B1 METABOLISM | 1.61E+00 | 2.34E-01 | 6 |
| 84 | METAPATHWAY BIOTRANSFORMATION PHASE I AND II | 1.56E+00 | 2.35E-01 | 115 |
| 85 | CYSTEINE AND METHIONINE CATABOLISM | 1.60E+00 | 2.40E-01 | 11 |
| 86 | ARYL HYDROCARBON RECEPTOR PATHWAY WP2586 | 1.56E+00 | 2.41E-01 | 42 |
| 87 | CYTOSOLIC DNASENSING PATHWAY | -1.47E+00 | 2.41E-01 | 52 |
| 88 | CANNABINOID RECEPTOR SIGNALING | 1.61E+00 | 2.42E-01 | 24 |
| 89 | ANTIVIRAL AND ANTIINFLAMMATORY EFFECTS OF NRF2 ON SARSCOV2 PATHWAY | 1.55E+00 | 2.44E-01 | 26 |
| 90 | OSTEOBLAST SIGNALING | -1.47E+00 | 2.45E-01 | 7 |
| 91 | PILOCYTIC ASTROCYTOMA | 1.56E+00 | 2.46E-01 | 6 |
| 92 | mRNA PROTEIN AND METABOLITE INDUCATION PATHWAY BY CYCLOSPORIN A | 1.61E+00 | 2.48E-01 | 7 |
| 93 | DISORDERS OF FOLATE METABOLISM AND TRANSPORT | -1.47E+00 | 2.48E-01 | 12 |
| 94 | SARSCOV2 REPLICATION ORGANELLE FORMATION | 1.57E+00 | 2.49E-01 | 6 |
| 95 | GLYCEROPHOSPHOLIPID BIOSYNTHETIC PATHWAY | 1.54E+00 | 2.49E-01 | 27 |

**Supplementary Figure S2. Table listing all significative enriched gene sets in treated BxPC3 cells.** Rankings based on FDR score. ± NES indicates upregulation or downregulation respectively of gene set in treated BxPC3 cells.

**Supplementary Figure S2**
