## Supplementary Figure 3 for "PGM3 inhibition Shows cooperative Effects With Erastin inducing Pancreatic cancer cell death via activation of the Unfolded Protein Response"

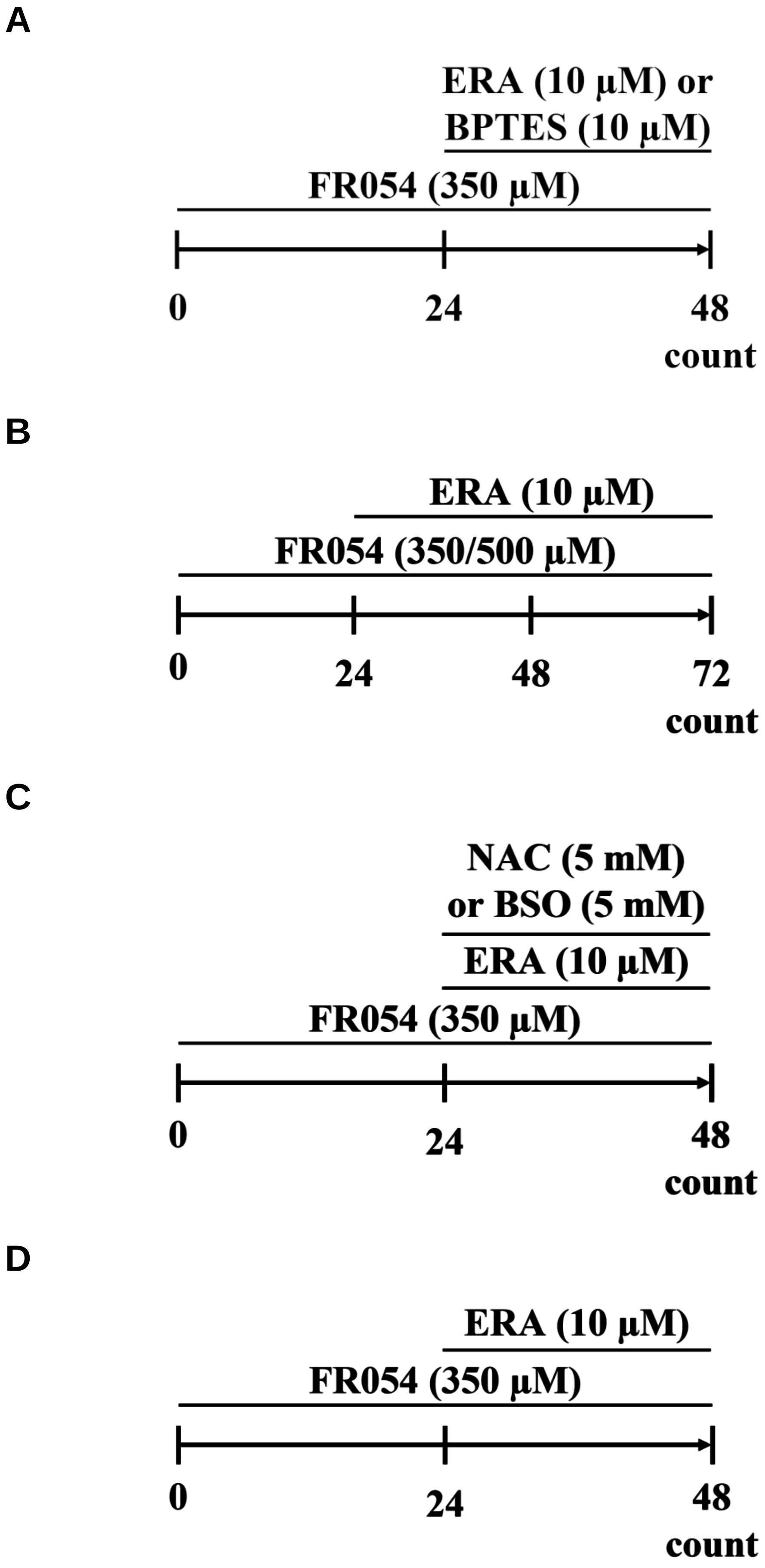


**Supplementary Figure S3. Experimental setting of the different treatments described in the main text.**

**Supplementary Figure S3**
