## Supplementary Figure 4 for "PGM3 inhibition Shows cooperative Effects With Erastin inducing Pancreatic cancer cell death via activation of the Unfolded Protein Response"

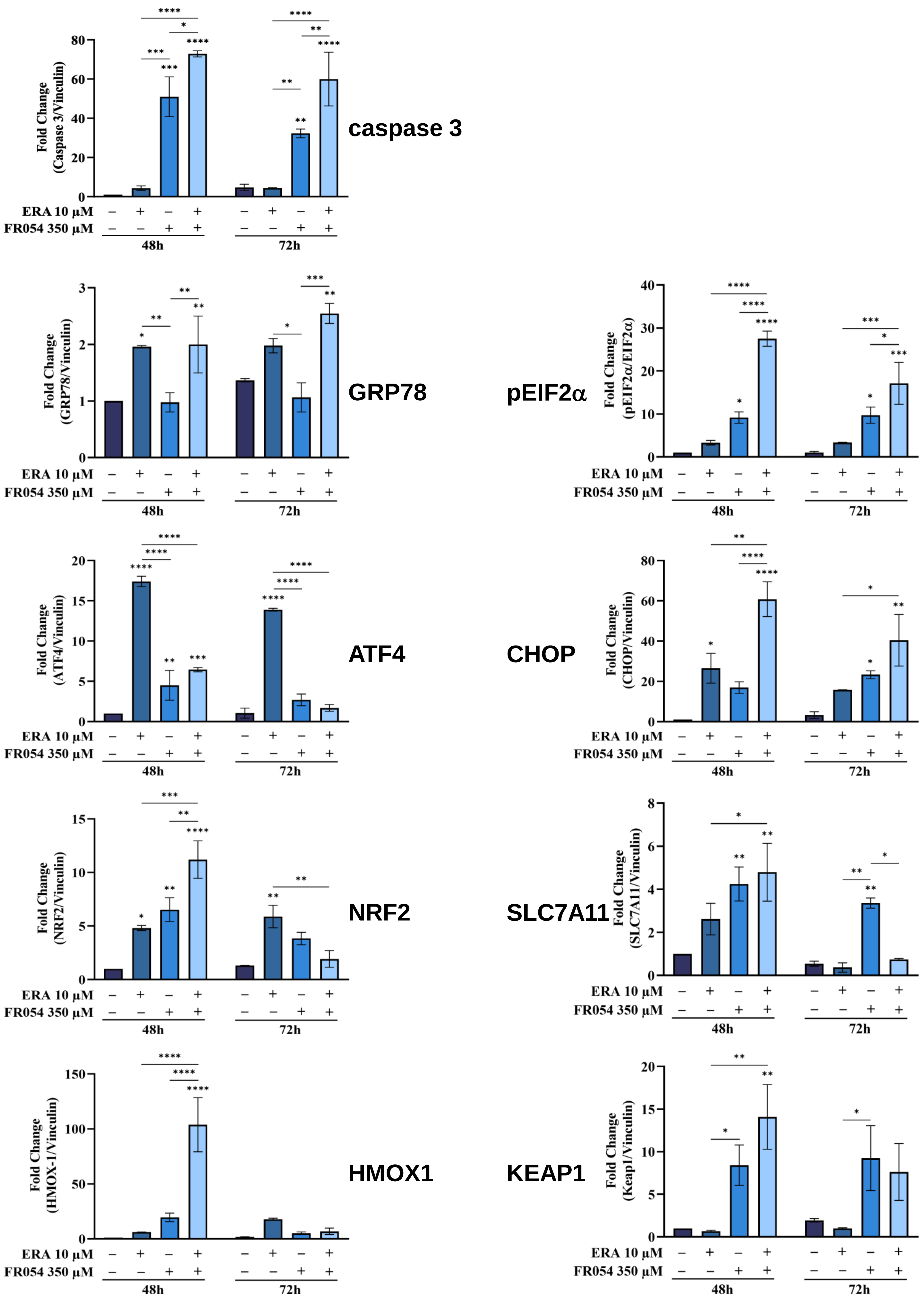


Supplementary Figure S4

**Supplementary Figure S4. The combined treatment of MiaPaca-2 cells with FR054 and erastin affects UPR and oxidative stress response proteins expression.** Densitometric analysis of immunoblot showing the effects of the two compounds on different UPR and oxidative stress response proteins evaluated at 48h and 72h post ERA, FR054 and their combination treatments. The immunoblot images are quantified by ImageJ software. *$p < 0.05$, **$p < 0.01$ ***$p < 0.001$, ****$p < 0.0001$. The data are presented as mean ± SD from three independent experiments.

**Supplementary Figure S4**
