## Supplementary Figure 6 for "PGM3 inhibition Shows cooperative Effects With Erastin inducing Pancreatic cancer cell death via activation of the Unfolded Protein Response"

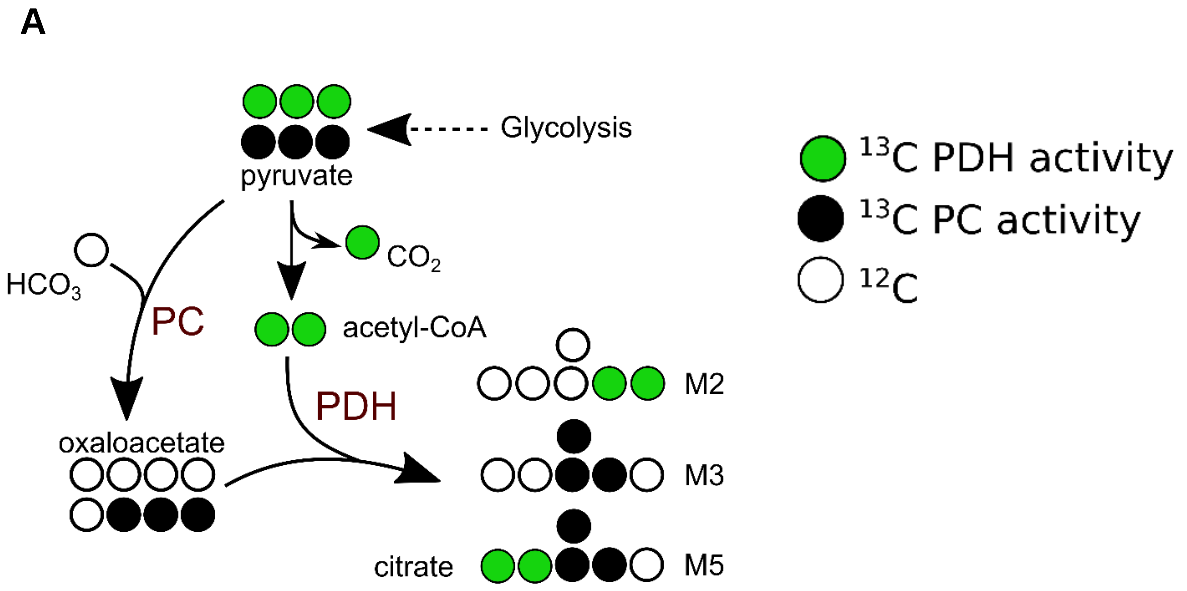


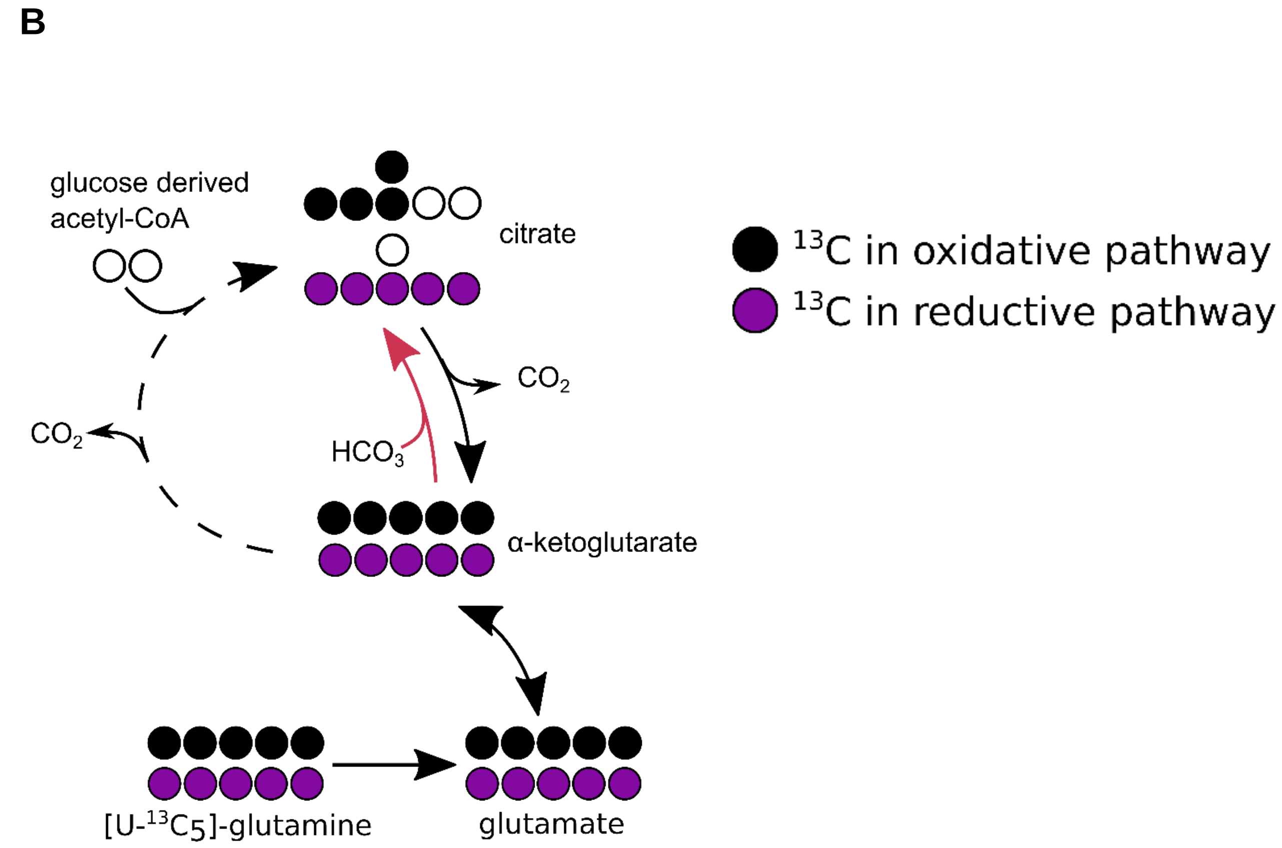


**Supplementary Figure S6. Isotopomers labeling for metabolomics study in MiaPaCa-2, BxPC3 and PANC1 cell lines.** (A) Picture describing pyruvate, citrate, α-ketoglutarate and glutamate isotopomers from [U-13]-glucose or (B) [U-13]-glutamine. For more information refer to main text.

**Supplementary Figure S6**
