## Supplementary Figure 7 for "PGM3 inhibition Shows cooperative Effects With Erastin inducing Pancreatic cancer cell death via activation of the Unfolded Protein Response"

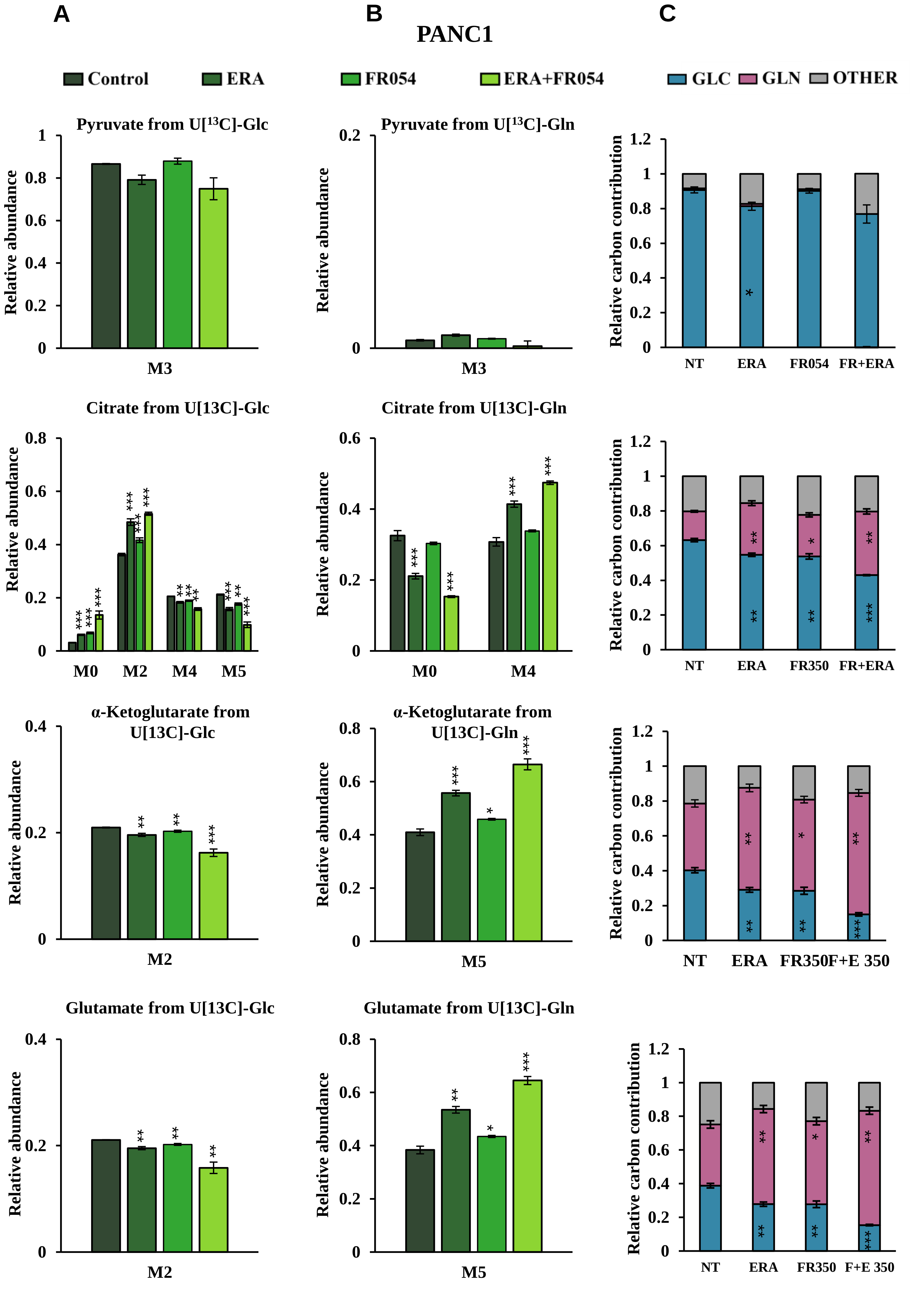


Supplementary Figure S7

**Supplementary Figure S7. Combined treatment in PANC1 cells favors glutaminolysis over glycolysis.** (A) and (B) 13C labeling of pyruvate, citrate, a-ketoglutarate and glutamate from PANC1 cells incubated in [U-$^{13}C_6$]-glucose or [U-13C5]-glutamine medium for 48h. (C) Relative carbon fractional contribution of glucose and glutamine to pyruvate, citrate, a-ketoglutarate and glutamate formation. *p < 0.05, **p < 0.01 ***p < 0.001. The data are presented as mean ± SEM from three independent experiments.

**Supplementary Figure S7**
